# Stretch-activated ion channels identified in the touch-sensitive structures of carnivorous Droseraceae plants

**DOI:** 10.1101/2020.12.15.422915

**Authors:** Carl Procko, Swetha E Murthy, William T Keenan, Seyed Ali Reza Mousavi, Tsegaye Dabi, Adam Coombs, Erik Procko, Lisa Baird, Ardem Patapoutian, Joanne Chory

## Abstract

In response to touch, some carnivorous plants such as the Venus flytrap have evolved spectacular movements to capture animals for nutrient acquisition. However, the molecules that confer this sensitivity remain unknown. We used comparative transcriptomics to show that expression of three genes encoding homologs of the MscS-Like (MSL) and OSCA/TMEM63 family of mechanosensitive ion channels are localized to touch-sensitive trigger hairs of Venus flytrap. We focus here on the candidate with the most enriched expression in trigger hairs, the MSL homolog FLYCATCHER1 (FLYC1). We show that *FLYC1* transcripts are localized to mechanosensory cells within the trigger hair, transfecting FLYC1 induces chloride-permeable stretch-activated currents in naïve cells, and transcripts coding for FLYC1 homologs are expressed in touch-sensing cells of Cape sundew, a related carnivorous plant of the Droseraceae family. Our data suggest that the mechanism of prey recognition in carnivorous Droseraceae evolved by co-opting ancestral mechanosensitive ion channels to sense touch.

## Introduction

How organisms evolve new forms and functions is a question of major interest. For example, whereas most plants respond slowly to mechanical stimuli—such as touch, gravity, or changes in turgor pressure—by altering growth, some plants have additionally evolved rapid touch-induced movements. These movements can be used to deter herbivory, as is seen with the sensitive plant (*Mimosa pudica*), or to physically capture and digest mobile animals, as occurs in carnivorous plants (Chehab et al., 2008). The best known carnivorous plant, *Dionaea muscipula* (Venus flytrap) of the Droseraceae family, is one such plant that has evolved a remarkable and rapid touch response to facilitate prey capture (**supplemental video 1**). How this predator evolved its complex leaf form and function for this purpose has long puzzled scientists; indeed, Charles Darwin proclaimed the species “one of the most wonderful [plants] in the world” (Darwin, 1875).

Several groups have described the events that evoke trap closure in Venus flytrap in detail. The plant leaf consists of an open bilobed trap, to which animal prey are attracted by volatile secretions (Kreuzwieser et al., 2015; Lloyd, 1942). There, the animal comes into contact with one of 3-4 mechanosensory trigger hairs on each lobe. Bending of any one trigger hair generates an action potential in sensory cells at the base of the trigger hair that propagates through the trap (Benolken & Jacobson, 1970; Burdon-Sanderson, 1873) and is correlated with an increase in cytosolic calcium in leaf trap cells (Suda et al., 2020). Two action potentials (or two touches) in a short period are required to likely further increase cytosolic calcium levels above a necessary threshold to initiate rapid closure of the trap, which ensnares the prey (Brown & Sharp, 1910; Suda et al., 2020). Trap closure occurs in as little as 100 milliseconds, representing one of the fastest movements in the plant kingdom (Forterre et al., 2005). These electromotive properties were first described in the 1870s (Burdon-Sanderson, 1873); however, almost a century and a half later, the molecular underpinnings of these events remain elusive.

Early recordings from Venus flytrap trigger hairs suggest that, based on the time scale of action potential generation, mechanosensitive ion channels may play a role in transducing force into electrical signals (Benolken & Jacobson, 1970; Jacobson, 1965). To identify possible ion channels required for the touch response in Venus flytrap, we sought to find genes that are preferentially expressed in trigger hairs.

## Results

The Venus flytrap genome is very large (Palfalvi et al., 2020)—approximately 20x the size of that of the commonly studied model plant *Arabidopsis thaliana* (**Figure 1-supplemental Table 1**)—and lacks available inbred strains. Therefore, we established a clonal propagation system for the collection of genetically identical material (**Figure 1A, B**, and **Figure1-figure supplement 1**). Using these clones, we generated a *de novo* transcriptome representing genes expressed in trap tissue from Illumina-based short sequencing reads. The transcriptome of our clonal strain consists of almost 28,000 unique transcripts with open reading frames coding for predicted proteins of at least 100 amino acids. This is larger than the approximately 21,000 genes previously predicted by genome sequencing (Palfalvi et al., 2020), reflecting the presence of multiple isoforms in our transcriptome, possible heterozygosity and strain differences, and/or differences in genome/transcriptome completeness, indicated by BUSCO analysis (see Materials and Methods). When we compared the transcriptome of trigger hairs with that of only the leaf traps, 495 protein-coding genes were differentially-enriched by greater than 2-fold in the trigger hair, whereas 1,844 were similarly enriched in the trap (**Figure 1-supplemental Tables 2** and **3**). Based on homology to *Arabidopsis*, many genes preferentially expressed in leaf traps were associated with photosynthetic function, whereas those more highly expressed in the trigger hairs included transcription factors and genes that may affect cellular and organ structure (Figure 1-supplemental Tables 4 and **5**).

**Figure 1.**
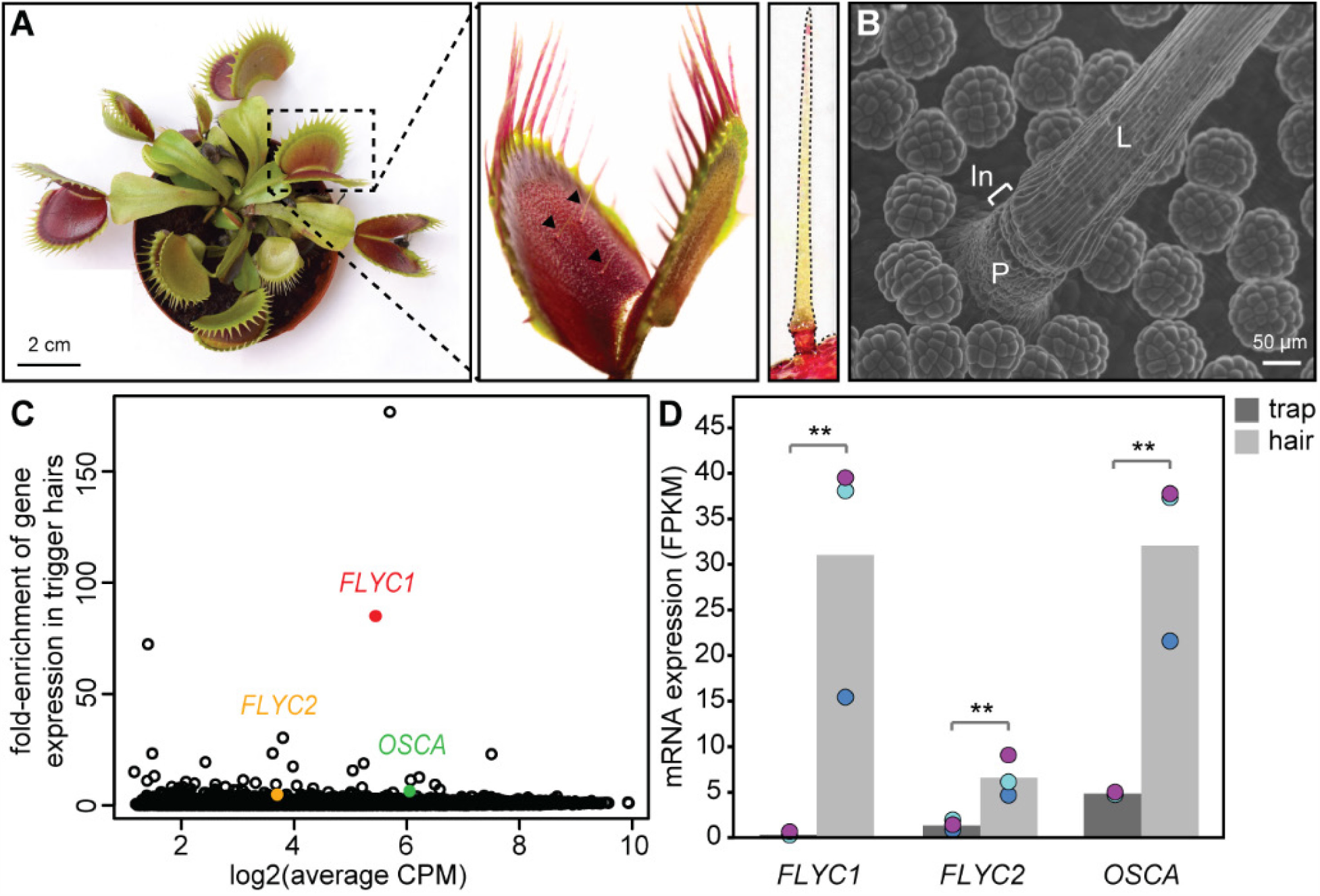
Identification of putative mechanosensory channels in the Venus flytrap trigger hair. (**A**) Representative image of a soil-grown Venus flytrap clone (left), Venus flytrap leaf (center), and single trigger hair (right). Black arrowheads in center picture indicate trigger hairs on leaf. (**B**) Scanning electron micrograph of a trigger hair. Cells of the lever (L), indentation zone (In) and podium (P) are indicated. Also seen are the digestive glands on the floor of the lobe. (**C**) Fold-enrichment of protein-coding genes of >100 amino acids in length (black circles) in the trigger hair relative to the trap. *FLYC1, FLYC2* and *OSCA* are shown in red, orange and green, respectively. CPM, counts per million of mapped sequencing reads. (**D**) Average Fragments Per Kilobase of transcript per Million mapped reads (FPKM) for *FLYC1, FLYC2* and *OSCA* in traps and trigger hairs. Dots of the same color indicate paired biological replicates. **FDR < 0.005.

To find potential mechanosensitive ion channels, we screened our trigger hair-enriched transcriptome for transcripts coding for likely multi-pass transmembrane proteins. Of 45 such transcripts, three coded for possible candidates based on homology to *Arabidopsis* proteins. Two of these shared homology to MSL family proteins (transcript IDs comp20014_c0_seq1 and comp28902_c0_seq1), and one to the OSCA family (comp16046_c0_seq1) (Haswell & Meyerowitz, 2006; Murthy et al., 2018). We call these genes *FLYCATCHER1* and *FLYCATCHER2* (*FLYC1* and *FLYC2*), and *DmOSCA*, respectively. We did not observe enriched expression of homologs to the mechanically activated PIEZO channels. Strikingly, one of these MSL-related transcripts, *FLYC1*, was expressed 85-fold higher in trigger hairs than in trap tissue (**Figure 1C, D**), and was the second highest enriched gene (a putative terpene synthase was 176-fold enriched; see Materials and Methods). By contrast, the other two putative ion channels were less than 7-fold differentially expressed. To validate our transcriptome, we verified the transcript sequence from cDNA of *FLYC1*. In addition, we used Sanger sequencing of PCR products generated from a genomic DNA template to assemble the complete gene sequence. 32 single nucleotide polymorphisms (SNPs) were detected between the two *FLYC1* alleles of our clonal strain, of which only 2 were found in the coding region and were both silent. Similarly, when we compared our coding sequence to that of another sequenced strain (Palfalvi et al., 2020), we found an additional four SNPs, of which only one caused an amino acid change (**Figure 1-figure supplement 2**). The scarcity of SNPs causing amino acid changes when compared to the abundance found in introns is likely indicative of selection for the protein product.

*FLYC1* and *FLYC2* code for predicted proteins of 752 and 897 amino acids, respectively, with homology to *Arabidopsis* MSL10 and MSL5 (**Figure 2A**). Ten MSL (MscS-like) proteins have been identified in *Arabidopsis* based on their similarity to the bacterial <u>m</u>echano<u>s</u>ensitive <u>c</u>hannel of <u>s</u>mall conductance, MscS (Haswell & Meyerowitz, 2006). In *Escherichia coli*, MscS opens to allow ion release upon osmotic down-shock and cell swelling, thereby preventing cell rupture (Booth & Blount, 2012; Levina, 1999). In *Arabidopsis*, MSL8 functions in pollen rehydration, whereas the roles of the remaining MSLs in mechanosensory physiological processes is unclear (Hamilton et al., 2015). Notably, MSL10 forms a functional mechanosensitive ion channel with slight preference for chloride when heterologously expressed in *Xenopus laevis* oocytes (Maksaev & Haswell, 2012). Furthermore, along with other members of the MSL family, MSL10 accounts for stretch-activated currents recorded from root protoplasts (Haswell et al., 2008). Whereas FLYC1 and FLYC2 are 38.8% identical (**Figure 2-figure supplement 1**), they share 47.5% and 35.5% identity with MSL10 respectively, with most variation in the cytoplasmic N-terminus. Compared to MscS, FLYC1 and FLYC2 have three additional predicted transmembrane helices, six in total, with the C-terminus of the proteins sharing highest homology (**Figure 2B, C**).

**Figure 2.**
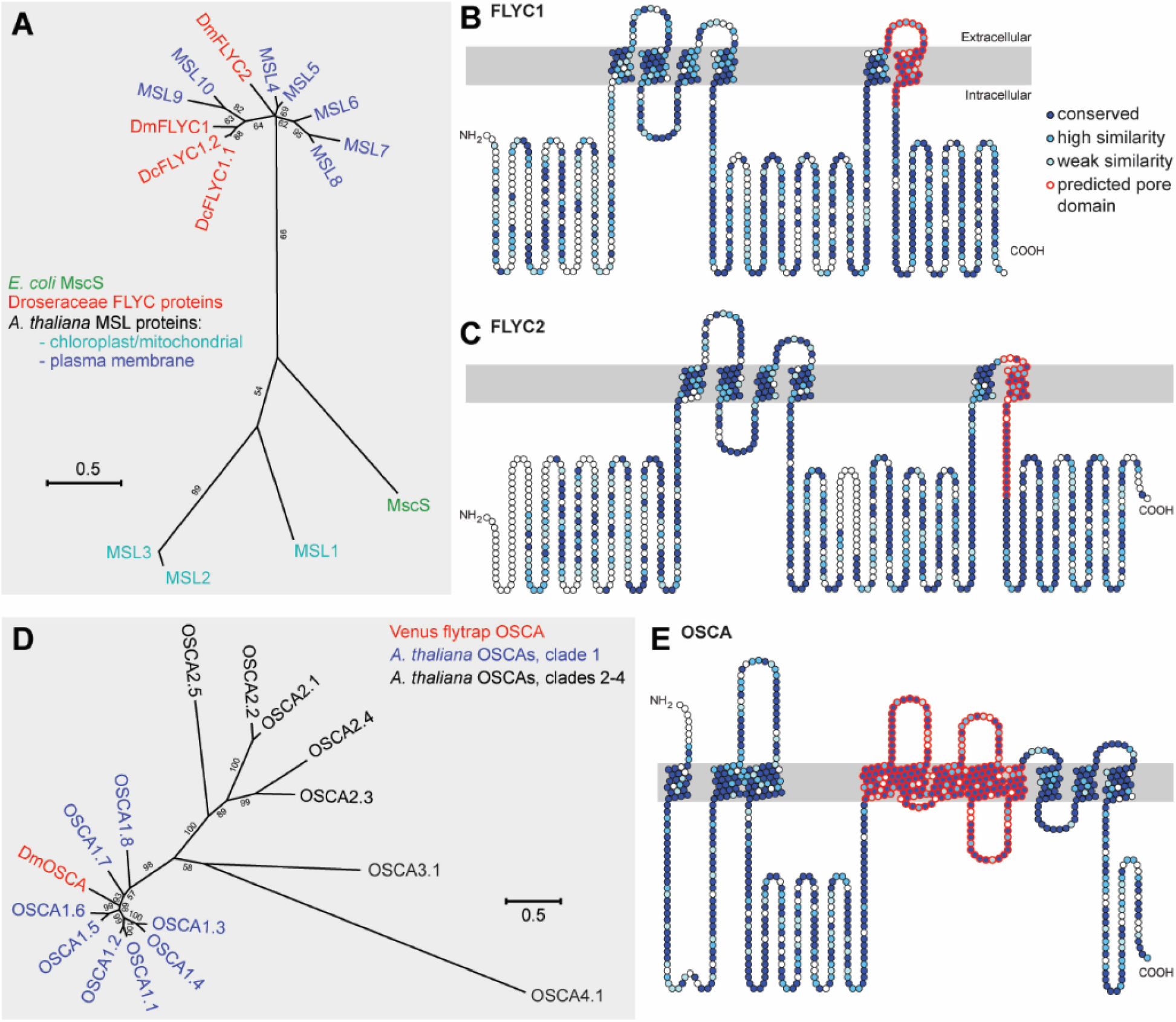
Molecular phylogenetic relationship of FLYCATCHER and OSCA proteins. (**A**) Phylogenetic analysis by maximum likelihood method to show the relationship between the conserved MscS domain (Haswell & Meyerowitz, 2006) of *Escherichia coli* MscS protein, *Arabidopsis thaliana* MSL proteins (MSL1-MSL10), and *Dionaea muscipula*/Venus flytrap (DmFLYC1 and DmFLYC2) and *Drosera capensis*/Cape sundew (DcFLYC1.1 and DcFLYC1.2) FLYCATCHER proteins. (**B, C**) Predicted topology for DmFLYC1 (**B**) and DmFLYC2 (**C**) proteins. (**D**) Phylogenetic analysis by maximum likelihood method showing the relationship between *Arabidopsis thaliana* OSCA family proteins and Venus flytrap DmOSCA. (**E**) Predicted topology for DmOSCA protein. In **A** and **D**, bootstrap values > 50 are shown; scale, substitutions per site. In **B, C** and **E**, amino acid residues that are conserved with *Arabidopsis* MSL10, MSL5 and OSCA1.5, respectively, are indicated in dark blue circles, whereas residues similar in identity are indicated in lighter blue circles. The predicted pore domain for each protein is indicated in red circles.

*DmOSCA* encodes a predicted 754 amino acid protein with highest homology to *Arabidopsis* OSCA1.5 (64% identity), which belong to a 15 member family of OSCA proteins in *Arabidopsis* (**Figure 2D, E**). Although initial identification characterized the OSCA family as hyperosmolarity-activated calcium channels, it has been demonstrated that several members of the OSCA family are mechanosensitive ion channels that are non-selective for cations with some chloride permeability (Murthy et al., 2018; Yuan et al., 2014). Fly and mammalian orthologues, Tmem63, also encode mechanosensitive ion channels, suggesting that the molecular function of the OSCA genes is conserved. Furthermore, purification and reconstitution of AtOSCA1.2 in proteoliposomes induce stretch-activated currents, indicating that these proteins are inherently mechanosensitive (Murthy et al., 2018). Notably, mutant OSCA1.1 plants have stunted leaf and root growth when exposed to hyperosmotic stress (Yuan et al., 2014), possibly as a consequence of impaired mechanotransduction in response to changes in cell size. This suggests an ancestral role for these channels as osmosensors, similar to the MSL family.

We sought to find which cells in the trigger hair express *FLYC1, FLYC2*, and *DmOSCA*. The trigger hair can be divided into two main regions: a cutinized lever and a podium on which the lever sits (**Figure 1B**). An indentation zone at the top of the podium separates the two regions (**Figure 3 A, C**), and is where most flexure of the trigger hair occurs (Lloyd, 1942). Electrophysiological recordings have demonstrated that mechanical stimulation of the trigger hair generates action potentials from a single layer of sensory cells at this indentation zone, and not from other cells of the podium or lever (Benolken & Jacobson, 1970). Remarkably, by fluorescent *in situ* hybridization, we detected *FLYC1* transcript specifically in indentation zone sensory cells (**Figure 3C, B**, and **D**). No transcript was observed in cells of the lever or lower podium (**Figure 3B, E**). In contrast, *EF1α* transcript, a housekeeping gene, was detected throughout the trigger hair and trap (**Figure 3-figure supplement 1**). *FLYC1*-sense probes produced no signal above background (**Figure 3B**). Using similar *in situ* hybridization methods, we were unable to detect *FLYC2* transcripts, whereas *DmOSCA* was found at high levels within trigger hair sensory cells, but also low levels in other cell types (**Figure 3-figure supplement 2**). These results are consistent with our RNA-seq findings from whole trigger hairs, where *FLYC2* and *DmOSCA* were less enriched over background trap tissue compared to *FLYC1*, and in the case of *FLYC2* only weakly expressed (**Figure 1D**). Regardless, the expression profile of *FLYC1* and *DmOSCA* is consistent with them being at the site of touch-induced initiation of action potentials.

**Figure 3.**
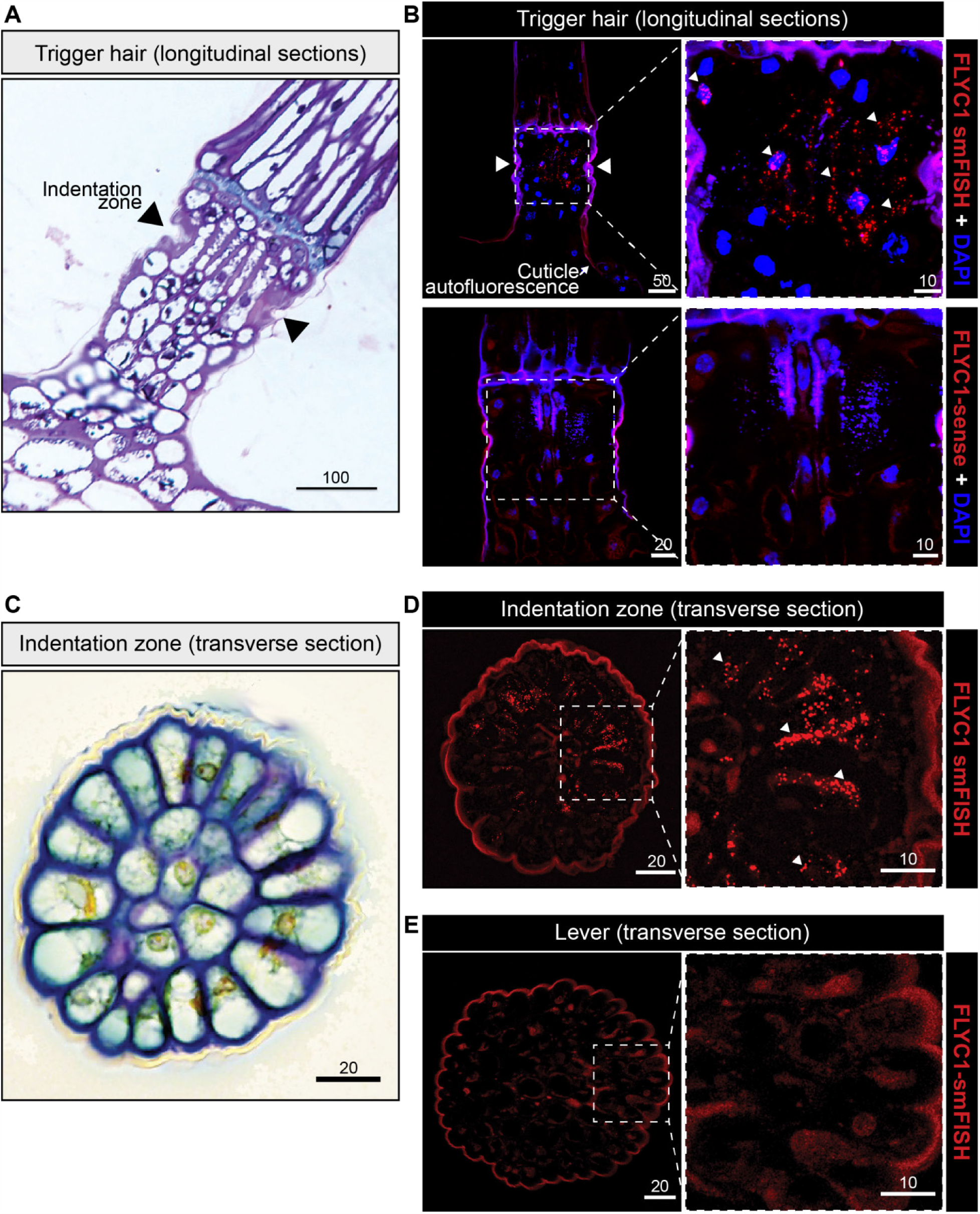
*FLYC1* mRNA localization in Venus flytrap trigger hairs. (**A**) Toluidine blue-stained longitudinal section through the base of a trigger hair. Elongated sensory cells are visible at the indentation zone (arrowheads). (**B**) Max projection through a longitudinal section after fluorescent *in situ* hybridization. (Top) *FLYC1* transcript (red) and DAPI (blue) at low (left) and high (right) magnification of the indentation zone. Note localization of the *FLYC1* transcript puncta in sensory cells. (Bottom) No signal was observed when using *FLYC1-*sense probes. High background fluorescence is observed in all channels. (**C**) Toluidine blue-stained transverse section through the indentation zone. The cells forming the outer ring are presumed to be the mechanosensors (Benolken & Jacobson, 1970). (**D**) Max projection through a transverse section after fluorescent *in situ* hybridization. *FLYC1* transcript (red) at low (left) and high (right) magnification. Note localization of the *FLYC1* transcript puncta predominately in the sensory cells of the indentation zone. (**E**) Max projection through a transverse section of the trigger hair lever. No transcript was observed. Scale bars, μm.

To test whether *FLYC1, FLYC2, and DmOSCA* were indeed mechanically-activated ion channels and could confer mechanosensitivity to naïve cells, we expressed human codon-optimized sequences of these genes in mechanically-insensitive HEK-P1KO cells (Dubin et al., 2017). Robust stretch-activated currents were recorded from FLYC1-expressing cells when negative pressure was applied to the recording pipette in the cell-attached patch configuration (**Figure 4, A, B**), but not from FLYC2-, or DmOSCA-expressing cells (**Figure 4-figure supplement 1**). Notably, overexpression of human codon-optimized MSL10 and of OSCA1.5 subcloned from *Arabidopsis thaliana* also failed to produce stretch-activated currents in our system (**Figure 4-figure supplement 1**). However, previous studies have shown that expression of MSL10 in oocytes does elicit stretch-activated currents, suggesting that trafficking of these plant proteins is largely compromised in mammalian cells (Maksaev & Haswell, 2012). Given the high transcript enrichment, localization in sensory cells, and mechanosensitivity, we focused on FLYC1 as a likely functional mechanosensor in Venus flytrap sensory hairs.

**Figure 4.**
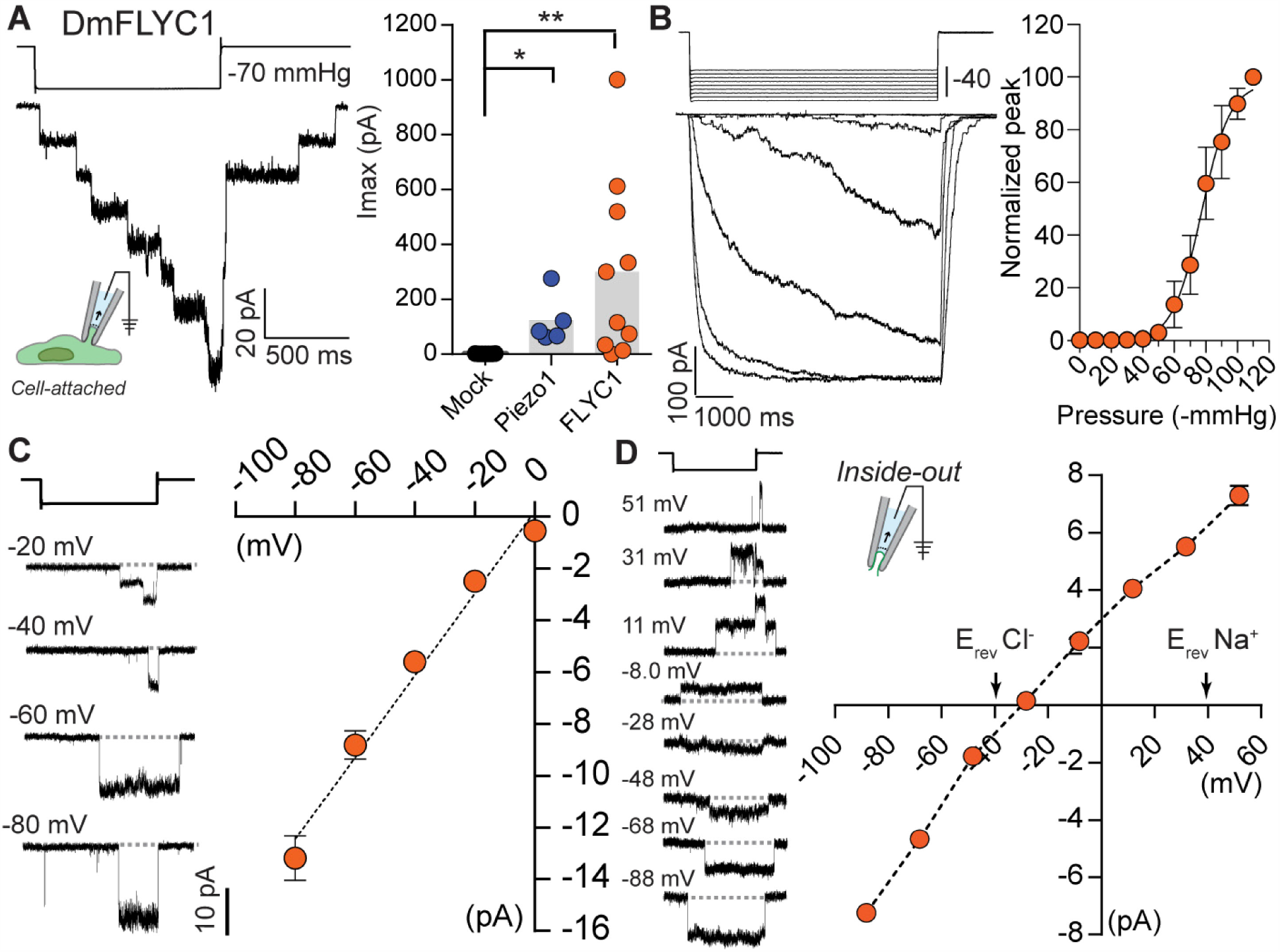
FLYC1 induces stretch-activated currents. (**A**) Left, representative trace of stretch-activated current recorded from FLYC1 expressing HEK-P1KO cells in the cell-attached patch clamp configuration at -80mV membrane potential in response to -70mmHg pipette pressure. Stimulus trace illustrated above the current trace. Right, quantification of maximal current response from cells transfected with mock (N=7), mouse *PIEZO1* (N=5), or *FLYC1* plasmid (N=10). *p*=0.0251 (Mock vs. *PIEZO1*); *p*=0.0070 (Mock vs. *FLYC1*); Dunn’s multiple comparison test. (**B**) Left, currents in response to graded negative pressure steps from 50 to 90 mmHg (Δ 5 mmHg) at -60 mV membrane potential. Right, average pressure-response curve normalized to peak current across cells (pressure-response curve for absolute peak values is plotted in Figure 4-figure supplement 2). A fit with Boltzman equation revealed P_50_ value of 77 mmHg (N=9). (**C**) Left, representative single channel traces in response to stretch at the indicated membrane potential. Currents in A-C were recorded in pipette solution containing (in mM) 130 NaCl, 5 KCl, 1 CaCl_2_, 1 MgCl_2_, 10 TEA-Cl and 10 HEPES (pH 7.3), and the bath solution containing 140 KCl, 1 MgCl_2_, 10 glucose and 10 HEPES (pH 7.3). Right, average I-V relationship of stretch-activated single channel currents from *FLYC1* transfected cells (N=6). (**D**) Left, representative stretch-activated single channel currents recorded from excised inside-out patch configuration in asymmetrical NaCl solution at the indicated membrane potential. Right, Average I-V of stretch-activated single channel currents in asymmetrical NaCl solution (E_rev_: -30.0 ± 1.4 mV (N=7)). Extracellular solution (pipette solution) composition was (in mM) 150 NaCl and 10 HEPES (pH 7.3) and intracellular solution (bath solution) composition was 30 NaCl, 10 HEPES and 225 Sucrose (pH 7.3).

Further characterization of FLYC1 channel properties indicated that the pressure required for half maximal activation (P_50_) of FLYC1 was 77.3 ± 4.0 mmHg (N=9) (**Figure 4B**), the channel had a conductance of 164 ± 9 pS (N=6) (**Figure 4C**), and upon removal of the stretch stimulus, the currents decay with a time constant of 167 ± 34 ms (N=5). In land plants and green algae, chloride-permeable channel opening is associated with membrane depolarization, due to the efflux of chloride ions down their electrochemical gradient (Beilby, 2007; Hedrich, 2012). Therefore, we tested whether FLYC1 is permeable to chloride by recording stretch-activated FLYC1 currents from excised patches in asymmetrical NaCl solution. FLYC1 exhibited a preference for chloride over sodium with a P_Cl_/P_Na_ ratio of 9.8 ± 1.8 (N=7) (**Figure 4D** and **Figure 4-figure supplement 2B**).

Bacterial MscS are bona fide mechanically activated ion channels, with in-depth structural and molecular understanding of how they are gated by membrane tension (Booth & Blount, 2012). Much less is known about the structure and function of plant MSLs. Although homology modelling of MSL10 with MscS has hinted at certain residues that alter channel conductance, molecular determinants of selectivity and gating in MSL10 remain largely unknown (Maksaev & Haswell, 2013; Maksaev et al., 2018). Intrigued by the high sequence similarity in the C-terminus of MscS, MSL10, and FLYC1 proteins (**Figure 4-figure supplement 3**), we asked to what extent the molecular architecture of the putative pore-lining helix is shared among these channels. Homology-modelling of FLYC1 with the known MscS structure (Bass et al., 2002) suggested that certain features of the pore-forming transmembrane domain (TM6) are indeed conserved. These include hydrophobic residues within the part of the helix that lines the pore (TM6a), and a glycine kink at G575 followed by an amphipathic helix (TM6b) that lies parallel to the inner plasma membrane surface (**Figure 4-figure supplement 3**). Based on our result that FLYC1 has a high preference to chloride we tested whether the positively-charged lysines on either side of the putative pore (K558 and K579) may confer pore properties. We substituted lysines at position 558 and 579 with glutamate and measured P_Cl_/P_Na_. Whereas selectivity for chloride in both mutants remained unchanged, the K579E mutant exhibited smaller single channel currents at positive membrane potentials (**Figure 4-figure supplement 3**), suggesting that K579 is indeed in the vicinity of the pore. This analysis confirms that the predicted TM6 of FLYC1 is part of the pore-lining region of the channel, consistent with mutagenesis results in MSL10 (Maksaev et al., 2018). These observations are further supported by the cryo-EM (electron microscopy) structure of AtMSL1, which indicates a similar architecture of the last TM of the channel (Deng et al., 2020). Future structural studies on DmFLYC1 will better resolve how selectivity is determined in these channels.

If FLYC1 is indeed important for touch-induced prey recognition, we reasoned that its expression and function in mechanosensory structures is likely to be conserved across carnivorous Droseraceae plants. To test this, we investigated the largest genus in the family, *Drosera*, which includes ∼200 species of sundew (Poppinga et al., 2013). Sundews are characterized by touch-sensitive projections on their leaf surface called tentacles (**Figure 5A**) (Darwin, 1875; Lloyd, 1942; Poppinga et al., 2013; S. E. Williams, 1976). These tentacles typically secrete a glob of sticky mucilage from their head, which acts as a trapping adhesive when contacted by insect prey. Movement by the adhered insect results in action potentials along the tentacle (Williams & Pickard, 1972), often accompanied by radial movement of the tentacle towards the leaf center (Darwin, 1875). This traps the struggling insect against even more mucilage-secreting tentacles, allowing for digestion to occur (**Figure 5B**, and **supplemental videos 2** and **3**).

**Figure 5.**
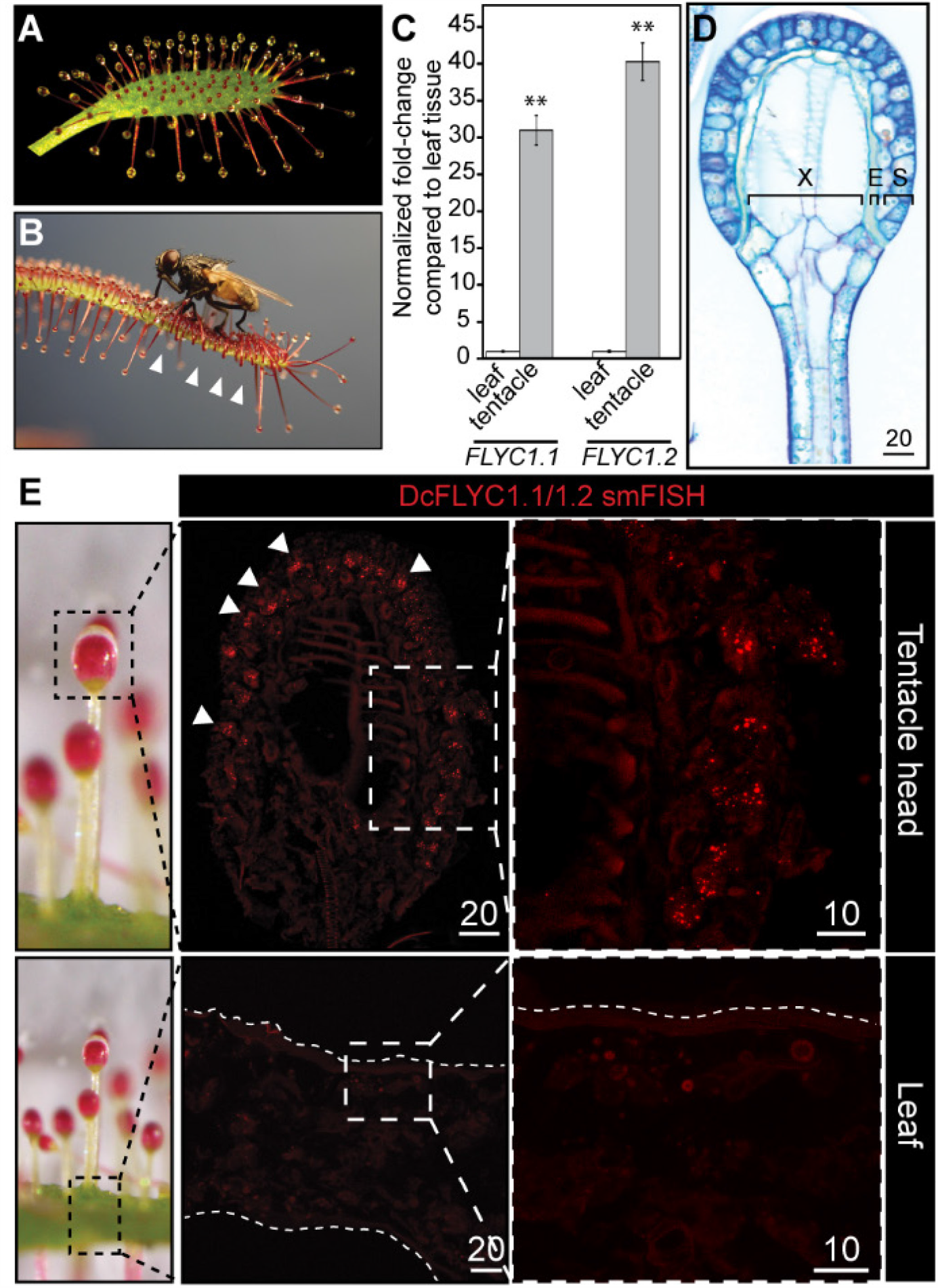
*DcFLYC1*.*1* and *DcFLYC1*. *2* localize to touch-sensitive structures of *Drosera*. (**A**) The Cape sundew leaf, showing tentacle projections with mucilage secretions. (**B**) Image of tentacle bending in response to insect (house fly) touch. Arrowheads mark examples of tentacles that have bent inward. (**C**) Relative expression of *DcFLYC1*.*1* and *DcFLYC1*.*2* in sundew tentacles versus tentacle-less leaves by qRT-PCR. \*\**p* < 0.005, moderated t test, results from four biological replicates of each tissue type. (**D**) Toluidine blue-stained longitudinal section through the head and upper neck of a Cape sundew tentacle. The head is composed of xylem (X), an endodermis-like layer (E), and secretory cells (S). (**E**) Max projection image, at low (left) and high magnification (right), showing collective localization of *DcFLYC1*.*1* and *DcFLYC1*.*2* mRNAs to the outer secretory cells of the tentacle head (top). Arrows indicate example secretory cells with *DcFLYC* puncta. No signal above background was observed in the leaf (bottom). Scale bar, µm.

We identified the expression of two *FLYC1* homologs in Cape sundew (*Drosera capensis*) by quantitative reverse transcriptase-(qRT-) PCR (Butts et al., 2016). Consistent with our findings from Venus flytrap, these two transcripts had 30- to 40-fold higher expression in tentacles compared to tentacle-less leaf tissue (**Figure 5C**). The cloned cDNAs of these two genes—which we call *DcFLYC1*.*1* and *DcFLYC1*.*2*—code for almost identical predicted protein products with 96.4% identity. They share 66.2% and 65.6% identity to Venus flytrap FLYC1, respectively (**Figure 2A**).

Cape sundew tentacles display variation in length depending on their position on the leaf (**Figure 5-figure supplement 1**), but share a similar mechanosensory head structure, with two outer layers of secretory cells (**Figure 5D**) (Lloyd, 1942; Williams & Pickard, 1972, 1974). The outermost secretory cells have previously been hypothesized to also be the site of touch sensation (Lloyd, 1942). Not only are these cells directly exposed to the mucilage environment on which the insect prey adheres and pulls, but they display a unique morphology of outer cell wall buttresses and plasma membrane crenellations (**Figure 5-figure supplement 2**) (Lloyd, 1942). In addition, cellulose fibrils extend from the outer cell wall into the cuticle in much the same way as they do in the Venus flytrap indentation zone (Sievers, 1968; Williams & Pickard, 1974). Strikingly, smFISH probes against *DcFLYC1*.*1* and *DcFLYC1*.*2* transcripts localized to the outer secretory cells, whereas no transcripts were observed in the leaf at the base of the tentacles (**Figure 5E**).

Heterologous expression of human codon-optimized *DcFLYC1*.*1* and *DcFLYC1*.*2* cDNA, independently or together in HEK-P1KO cells, did not result in stretch-activated currents (**Figure 5-figure supplement 3**). Similar to our findings with DmFLYC2 and DmOSCA, we speculate that the lack of activity could be due to incorrect folding and trafficking of these proteins. Nonetheless, our expression data is consistent with a conserved role for FLYC1 in two divergent species in carnivorous Droseraceae plants.

## Discussion

Here, we identify members of the FLYC and OSCA families of ion channels as candidate mechanosensors in rapid touch sensation in carnivorous plants. First, members of both families have enriched mRNA transcript expression in the sensory trigger hairs of Venus flytrap. Among these, *FLYC1* is a prime candidate as its mRNA is massively enriched in the putative mechanosensory cells that initiate transduction of the touch-induced signal. Second, FLYC1 forms a mechanically activated ion channel with properties that would facilitate generation of action potentials in sensory cells. Third, expression of *FLYC* genes is remarkably conserved in two morphologically disparate touch-sensitive structures from different genera in the Droseraceae family. In addition to the three candidates presented here, our results do not exclude the possibility that other mechanosensitive proteins could also contribute to the events that enable touch-induced prey capture. In the future, knockout experiments will be critical to determine which candidate mechanosensor(s) are absolutely necessary for functional prey detection and trap closure.

Using a mammalian cell-expression system, we detected robust macroscopic as well as single channel stretch-activated currents in FLYC1-expressing cells and identified residues important for channel properties, which strongly suggests that FLYC1 form mechanosensitive ion channels. However, we were unable to detect currents from the other channels tested, including *Arabidopsis* MSL10, which has previously been shown to induce currents only in oocytes but not in mammalian cells. We hypothesize that these channels fail to fold and traffic correctly to the mammalian cell membrane, or, alternatively, that the activation threshold for these channels might be higher than that of *DmFLYC1* (Venus flytrap), technically precluding us from recording reliable stretch-activated currents. Future functional studies on these genes in different expression systems and purification and reconstitution of proteins in lipid vesicles will conclusively determine whether these genes encode inherently mechanosensitive ion channels.

It is interesting to compare FLYC1 channel properties to its homologs, MSL10 and MscS. FLYC1 single channel conductance is 164 ± 9 pS when measured from cell attached patches in HEK-P1KO cells, whereas MSL10 and MscS channel conductance from oocytes is 103 ± 3 pS and 218 ± 2 pS, respectively (Maksaev & Haswell, 2011). FLYC1 and MSL10 stretch-activated currents have slow opening and closing kinetics with lack of inactivation in the presence of sustained pressure, unlike MscS (Maksaev & Haswell, 2011). The half-maximal activation for FLYC1 is two-fold lower than its bacterial orthologs MscS (P_50_ 188 ± 31 mmHg; Akitake et al., 2005) and MscL (P_50_ 147.3 ± 4.3 mmHg; Doerner et al., 2012). However, we observed that similar to MSL10 recordings in oocytes (Maksaev & Haswell, 2012), the cell membrane in our recordings often ruptured before current saturation (saturated Po was measured in only 3/9 cells) resulting in our measurement of FLYC1 P_50_ likely being an underestimation. Finally, P_Cl_/P_Na_ ratio of FLYC1 is higher than ratios obtained for MSL10 (P_Cl_/P_Na_: 5.9) and MscS (P_Cl_/P_K_: 1.2-3.0) (Cox et al., 2014; Maksaev & Haswell, 2012). In depth functional and structural characterization of these channels (FLYC1, MSL10, and MscS) under similar expression systems and recording conditions will further determine absolute differences in their properties and the underlying molecular architecture that govern these properties.

Our characterization of FLYC1 as a chloride-permeable mechanically activated ion channel suggests a possible mechanism to elicit rapid touch-induced movements in Venus flytrap. Due to the high electronegativity of sensory cells (Benolken & Jacobson, 1970; Hodick & Sievers, 1988), combined with a concentration gradient that likely favors efflux (Higinbotham et al., 1967), a chloride-permeable channel like FLYC1 could contribute to membrane depolarization. This is consistent with the finding that increasing concentrations of extracellular chloride ions reduced or abolished the electrical response of Venus flytrap sensory cells to a mechanical stimulus (Jacobson, 1974). FLYC1-mediated depolarization, likely in combination with FLYC2 and OSCA, would then initiate action potentials that propagate into the leaf through plasmodesmata clustered on the basal side of the sensory cells (Williams & Mozingo, 1971), eliciting trap closure (**Figure 6A**). The role of a mechanosensitive calcium permeable ion channel, OSCA, is noteworthy here. Indeed, it has been described that trigger hair movement causes a rise in cytosolic calcium and that this signal propagates through the trap preceding its closure (Suda et al., 2020); however, the propagation signal most likely involves a mechanism independent of mechanotransduction. A similar chain of events would occur in the sundew tentacle: prey contact with the tentacle results in DcFLYC-induced action potentials, possibly in combination with OSCA or other mechanosensory proteins not identified here, that may propagate through plasmodesmata along the outer cell layers of the tentacle mechanosensory head and stalk, evoking tentacle bending (**Figure 6B**). A method for transformation of Venus flytrap has been recently reported (Suda et al., 2020). By generating targeted knockouts using this method, future studies may be able to address which or if all of these channels are required redundantly for touch responses in the plant. Nonetheless, our findings are consistent with FLYC1 being one of the bona fide sensors of touch in these “most wonderful” plants, and shed new light on how carnivorous plants evolved to sense touch by co-opting ancestral, osmosensing mechanosensitive channels.

**Figure 6.**
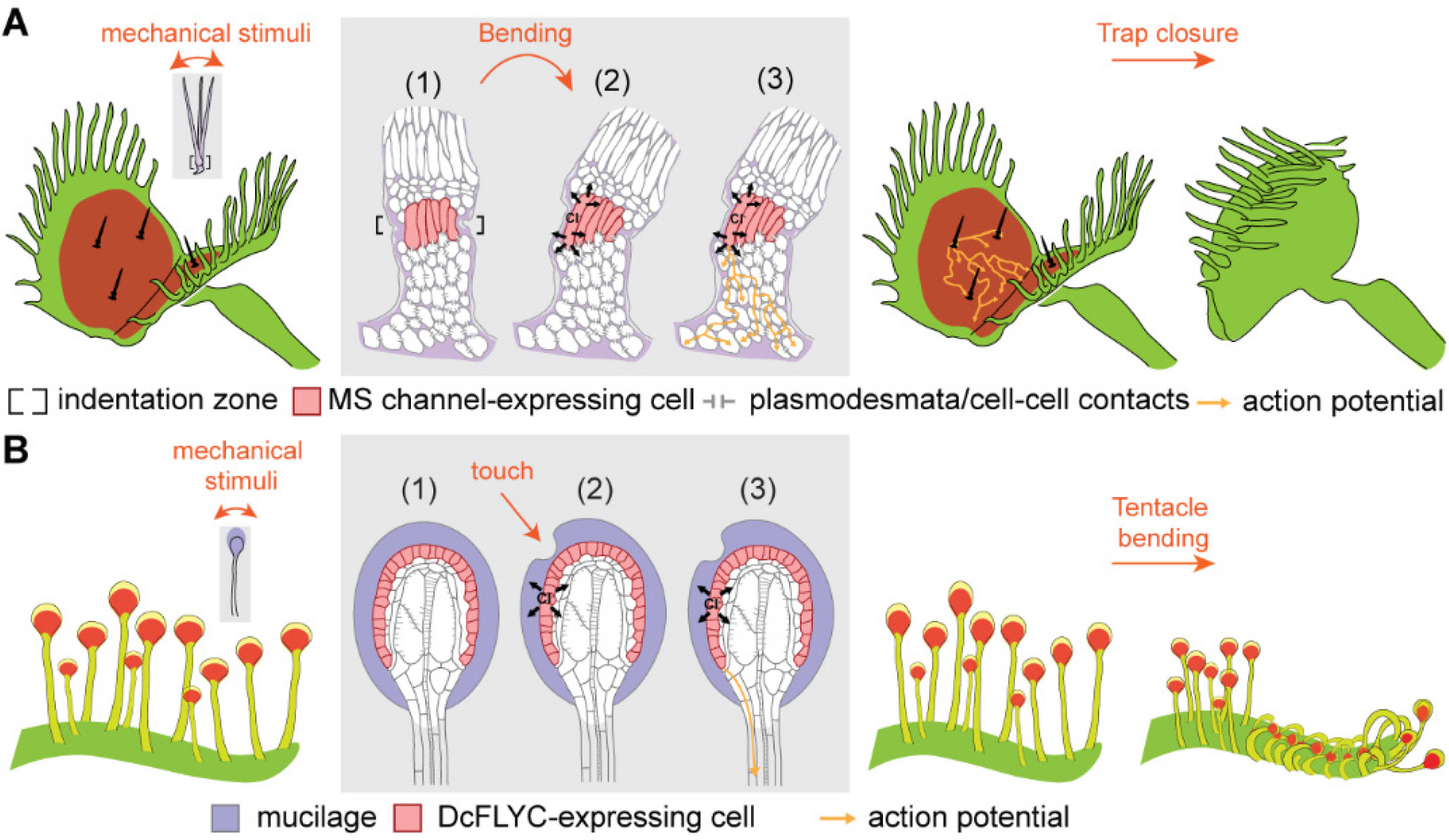
Model of touch-induced movements in carnivorous Droseraceae plants. (**A**) In Venus flytrap, mechanical stimulation of the trigger hair by a prey animal causes bending at the indentation zone sensory cells (1), leading to mechanically induced activation of mechanosensitive (MS) channels and chloride efflux (2). This triggers an action potential that propagates from the base of the sensory cells to cells of the podium via plasmodesmata (3) (Williams & Mozingo, 1971). Propagation of action potentials through the lobe of the leaf results in trap closure. (**B**) In sundew tentacles, the pulling of mucilage could lead to activation of DcFLYC1.1 and DcFLYC1.2 proteins in the outer cell layer of tentacle heads, triggering a propagating action potential down the tentacle stalk.

## Acknowledgements

We thank S. Morrison, K. Asahina, M. Stableford, S. McDowell, E. Perozo, and members of the Chory and Patapoutian laboratory for assistance and helpful discussions. We acknowledge Elizabeth Haswell for facilitating a collaboration between the Chory and Patapoutian laboratories.

## Author contributions

C.P., S.E.M., W.T.K., S.A.R.M., A.P., and J.C. conceived and designed experiments. C.P. performed gene expression analysis. S.E.M. recorded and analyzed electrophysiology data. W.T.K. performed and analyzed *in situ* expression. C.P., T.D., and S.A.R.M. cloned and generated *FLYC* cDNAs. A.C. generated FLYC1 mutants. E.P. performed FLYC1 homology modeling. L.B. performed light microscopy. C.P. and S.E.M. drafted the manuscript, and all authors contributed to finalization of the manuscript.

## Data and materials availability

Raw and processed data for the Venus flytrap transcriptome assembly and its analysis are available at the NCBI Sequencing Read Archive and Gene Expression Omnibus under accession numbers PRJNA530242 and GSE131340. Cloned cDNA sequences for *FLYC* genes are deposited at GenBank. All other data are available in the main text or supplementary materials.

## Materials and Methods

### Plant materials and growth conditions

Venus flytraps were propagated in tissue culture using methods modified from those previously described (Jang et al., 2003). Seeds (FlytrapStore.com, OR) were surface sterilized for 5 min using 70% ethanol with 0.1% Triton X-100; rinsed; and then seeded onto 1/3x Murashige and Skoog (MS) salts and vitamins (Caisson Labs), 3% sucrose, and 4.3 g/L gellan gum. Following germination, plants were selected for robust growth in tissue culture as well as large size to facilitate tissue manipulations. A single strain originating from a single seed (CP01) was chosen for further propagation by continual splitting of rhizomes. After 9-12 months in culture, the largest rosettes were transferred to soil (fine-grade sphagnum peat moss) and grown under greenhouse conditions (25-30°C; ∼16 h light 8 h dark, with overhead artificial red light added in evening hours to bring to 16 h light length). Soil was kept constantly moist using purified water. Plants were allowed to harden on soil for at least 2-3 months prior to experiments.

*Drosera capensis* var. rubra (Cape sundew) seeds (AK Carnivores, HI) were surface sterilized, germinated on plates, and seedlings transferred to soil and grown under greenhouse conditions.

### Imaging

For imaging of cell wall-stained tissue sections, freshly harvested plant tissue was dissected in 2% (v/v) paraformaldehyde and 1.25% (v/v) glutaraldehyde in 50 mmol/L PIPES buffer, pH 7.2 and fixed for 2 h in the same solution. Samples were dehydrated in a graded ethanol series and embedded in JB-4 Plus Embedding Media (Electron Microscopy Sciences) according to manufacturer’s instructions, with the exception that infiltration was performed at room temperature over 7 days. Some dehydrated samples were then stained with 0.1% eosin in 100% ethanol prior to embedding to help visualize the material during sectioning. Sections were cut at 4-6 μm and dried from a drop of dH2O onto Probe On Plus slides (ThermoFisher Scientific). Tissues sections were stained with 0.01% aqueous Toluidine Blue O, cover-slipped and examined with an Olympus BX51 microscope.

Scanning electron microscopy (SEM) was performed by imaging fresh plant tissue in the variable-pressure mode of a field emission-scanning electron microscope (Sigma VP; Zeiss) at 5 Pa of nitrogen with the variable-pressure secondary electron detector.

For time-lapse movies and light images of *D. capensis* during feeding, plants were fed with *Drosophila melanogaster* (Canton-S) or house flies (*Musca domestica*) and imaged using an Olympus Tough TG-5 camera. To aid in feeding, all insects were momentarily paralyzed by placing them in a tube on ice for 5-10 min immediately prior to plant feeding.

### Estimation of nuclear genome size

Nuclear genome size was estimated using flow cytometry methods similar to those reported previously (Arumuganathan & Earle, 1991), and were performed at Benaroya Research Institute, Seattle, WA. Nuclei from 50 mg of *Arabidopsis* leaf or Venus flytrap petiole tissue were suspended in 0.5 mL solution of 10 mM MgSO4, 50mM KCl, 5 mM Hepes pH 8.0, 3 mM dithiothreitol, 0.1 mg/mL propidium iodide (PI), 1.5 mg/mL DNAse free RNAse (Roche) and 0.25% Triton X-100. Nuclei were filtered through a 30 µm nylon mesh and incubated at 37°C for 30 min. Stained nuclei were analyzed with a FACScalibur flow cytometer (Becton-Dickinson). As an internal standard, samples included nuclei from chicken red blood cells (2.5 pg/2C). For each measurement, the PI fluorescence area signals (FL2-A) from 1000 nuclei were collected and analyzed and the mean positions of the G0/G1 peaks for the sample and internal standard were determined using CellQuest software (Becton-Dickinson). Nuclear DNA size estimates are an average of 4 measurements.

### Tissue collection for RNA-seq and qRT-PCR

To generate RNA-seq libraries from trigger hair tissue, trigger hairs were collected over the course of a month, 3-4 times a week during the hours of 6:30-9:30 pm under artificial red light. Only traps larger than 1.5 cm in length were used. 12 trigger hairs were dissected from the surface of two leaves at a time before snap-freezing in liquid N2 (< 5 min between dissection and freezing). Aliquots of trigger hairs were later pooled during RNA extraction. A sampling of traps (∼20) with the trigger hairs removed was collected simultaneously as comparison tissue. The experiment was performed in triplicate: 250 trigger hairs were collected for the first replicate, and 750 trigger hairs for each of replicates two and three.

For fed versus unfed trap samples, each tissue sample included two traps (> 1.5 cm in length), each fed with a single house fly for 24 h. Fed traps were opened at the time of harvest and the fly carcass removed before snap-freezing the tissue. Samples were collected in duplicate.

Cape sundew tentacles were harvested by removing fresh leaves from plants and placing these directly into liquid N2. While immersed in the liquid N2, the leaves were agitated and/or scraped to break off the tentacles. The tentacles and remaining leaf material were then separated for RNA extraction. Each sample type was collected in quadruplicate.

### RNA extraction for qRT-PCR and RNA-seq

Total RNA was extracted using a modified LiCl-based method similar to that described by others (Bemm et al., 2016; Jensen et al., 2015). Briefly, frozen plant material was ground to a powder and 700 μL of RNA extraction buffer added (2% CTAB, 2% polyvinylpyrrolidone K25, 100 mM Tris HCl pH 8.0, 25 mM Na-EDTA pH 8.0, 2M NaCl, 2% v/v β-mercaptoethanol). If necessary, tissue aliquots were pooled at this stage, or later during ethanol precipitation. Samples were vortexed 2 min and then incubated 65°C for 10 min, vortexing occasionally. Debris was removed by centrifugation and 600 uL chloroform was added to the supernatant and mixed. The sample was centrifuged at 10,000 rpm for 10 min. 1/3 vol 7.5 M LiCl was added to the aqueous phase which was then incubated overnight at 4°C with gentle mixing. RNA was pelleted by centrifugation at 10,000 rpm for 10 min at 4°C. The RNA pellet was washed with 70% ethanol, air-dried, and re-suspended in 100 μL H2O. The RNA was then re-precipitated by adding 0.1 vol 3M NaAcetate (pH 5.2), 2.5 vol 100% ethanol and an optional 1.5 μL GlycoBlue (Thermo Fisher) for low yield samples; mixing; and incubating for 1 h at -80°C. The RNA was pelleted by centrifugation at 12,000 rpm for 20 min at 4°C. The pellet was washed with 70% ethanol, air dried, and re-suspended in 30-50 uL H2O. These samples were then used directly to generate cDNA for qRT-PCR. Alternatively, samples for RNA-seq were re-suspended in 22 uL H2O, and to this was added 2.5 μL 10x TURBO buffer and 1 μL TURBO DNase (Thermo Fisher). Samples were incubated at 37°C for 20-30 min. 2.5 μL Inactivation Reagent was then added, and samples incubated for a further 5 min at room temperature, mixing occasionally. The resin was removed by centrifugation at 10,000 rpm for 1.5 min and transferring the supernatant to a new tube. RNA quality and yield was assessed by Agilent TapeStation before proceeding with sequencing library generation.

### RNA-Seq and Venus flytrap trap transcriptome assembly

Stranded mRNA-Seq libraries were prepared using Illumina TruSeq Stranded mRNA Library Prep Kit according to the manufacturer’s instructions. Libraries were quantified, pooled and sequenced at paired-end 125 bp reads using the Illumina HiSeq 2500 platform at the Salk NGS Core. Raw sequencing data was demultiplexed and converted into FASTQ files using CASAVA (v1.8.2). All samples were sequenced simultaneously on a single flow cell lane. The average sequencing depth was 8.8 million reads per library. Library quality was assessed using FastQC (v0.11.5) (Andrews, 2010) and Illumina adapter sequences and poor quality reads trimmed using Trimmomatic (0.36.0) with the suggested parameters by Trinity (Bolger et al., 2014).

Most aspects of our *de novo* transcriptome assembly and RNA-Seq analysis were performed using CyVerse cybercomputing infrastructure (Goff et al., 2011). So as to acquire as complete a representation of the Venus flytrap trap transcriptome as possible, we included reads collected from house fly-fed and unfed traps, in addition to our paired trigger hair and trigger hair-less trap samples to build our trap transcriptome (**Figure 1-supplemental Table 6**). We used Trinity (v2.5.1) for the *de novo* build (Grabherr et al., 2011), using default settings and an assembled contig length of greater than 300 nt. Our final transcriptome included 80,592 contigs, which were grouped into 77,539 components. In our manuscript and hereafter, each Trinity component is generally referred to as a “gene”. The maximum contig length was 15,873 nt; average and median lengths 867 and 536 nt, respectively; and N50 1,197 nt.

We used CEGMA (Core Eukaryotic Genes Mapping Approach) and BUSCO (Benchmarking Universal Single-Copy Orthologs) analyses to assess the completeness of our Venus flytrap trap transcriptome (Parra et al., 2007; Simao et al., 2015). Using CEGMA, of 248 ultra-conserved core eukaryotic genes tested for, all were present in our transcriptome, including 246 of which the coding sequences were defined as complete by CEGMA criteria. When we searched for the presence of near-universal single copy orthologs using BUSCO (v3.0; lineage: plantae; species: *Arabidopsis*), 94.7% of BUSCOs were present, 92.6% of which were defined as complete. This number is greater than that reported for the previously published Venus fly trap genome sequence (Palfalvi et al., 2020).

To further test the quality of our transcriptome, we aligned reads for each sample back to the transcriptome using Bowtie 2 (v2.2.4) (Langmead & Salzberg, 2012). For each sample, 80-90% of the paired reads mapped back concordantly, while the overall alignment rate was > 90%. These values were similar when the reads for each sample from a second sequencing run were independently aligned (see below).

TransDecoder (v1.0) (Haas et al., 2013) was used to find open reading frames (ORFs) that coded for possible proteins or incomplete protein fragments of at least 100 amino acids in length on the + strand. In total, 33,710 ORFs fulfilling these criteria were found (this number increased only a small amount, to 34,080, if the - strand was also included). Of these, 14,886 were identified as complete. Our values are not that dissimilar for a previously reported Venus flytrap transcriptome generated from multiple tissues (Jensen et al., 2015).

To assign homologous sequences from *Arabidopsis* to our protein-coding transcripts, we blasted all complete and partial polypeptides against the *Arabidopsis* TAIR10 proteome using default settings in Blastp (v2.2.29), with an e-value threshold of < 0.01 (Camacho et al., 2009). The top hit only was retained for downstream analysis. To predict the number of transmembrane passes per protein, we used TMHMM v. 2.0 (Krogh et al., 2001; Sonnhammer et al., 1998).

Our *de novo* Venus flytrap trap transcriptome is available through the National Center for Biotechnology Information (NCBI) Transcriptome Shotgun Assembly (TSA) database with accession number GHJF00000000. The uploaded transcriptome has the following modifications from that described above and used for differential gene expression analysis below: the last 42 nt of contig comp11005_c0_seq1 and the first 21 nt of comp46326_c0_seq1 were removed (possible adapter sequences), and 18 contigs were flagged as possible contaminants and deleted. The uploaded transcriptome includes 80,574 contigs.

### RNA-seq differential gene expression analysis

We used methods supported by Trinity (Haas et al., 2013) for finding genes differentially expressed between trigger hair and trigger hair-less trap tissue samples. To increase the sequencing read count number over any given contig, we performed a second sequencing run of all samples (SRA accession numbers SRR8834210, SRR8834209, SRR8834208, SRR8834207, SRR8834212 and SRR8834211). We trimmed the concatenated reads from both sequencing runs using Trimmomatic as described above, and aligned these to our *de novo* Venus flytrap trap transcriptome using Bowtie (Langmead et al., 2009). RSEM (v1.2.12) was used to find expected gene counts (Li & Dewey, 2011), and edgeR to find differentially-expressed genes using a paired experimental design for statistical testing (Robinson et al., 2010). For our analysis in edgeR (v3.12.1), we first filtered our gene list to include only those Trinity components whose transcript(s) included an ORF coding for a protein (or fragment thereof) of at least 100 amino acids in length. For components with more than one such protein assigned to them, the longest ORF was assumed to be the most relevant, and was used as the basis for assigning an *Arabidopsis* homolog to the gene/component. Genes that had counts per million (CPM) < 2 in over half the samples were excluded from the analysis. A table of genes passing these criteria showing gene expression values as mean CPM in traps and trigger hairs, as well as differential expression in trigger hairs versus traps calculated using edgeR algorithms with false discovery rate (FDR) can be found at the NCBI Gene Expression Omnibus (GEO) database (Edgar, 2002) with GEO Series accession number GSE131340 (https://www.ncbi.nlm.nih.gov/geo/query/acc.cgi?acc=GSE131340). This repository also includes a list of all protein-coding genes and blast results against the *Arabidopsis* proteome. Genes that had a fold-enrichment difference of 2-fold or more, in addition to FDR < 0.05, were designated as having trigger hair- or trap-enriched expression. These are shown in **Figure 1-supplemental Tables 2** and **3**. Note that fold-enrichment is calculated using edgeR algorithms, and not directly from mean CPM values. For gene ontology (GO) analysis (Figure 1-**supplemental Tables 4** and **5**), GO terms were assigned to each gene based on its homologous *Arabidopsis* protein (*Arabidopsis* TAIR10 annotation data downloaded April 2018). GO enrichment was performed using BiNGO 3.03 (biological process terms only) with Benjamini and Hochberg corrected p value < 0.05 (Maere et al., 2005).

The highest differentially-expressed gene in our trigger hair transcriptome coded for a protein with homology to a terpene synthase (transcript ID comp18811_c0_seq1; approximately 175-fold enrichment). It is possible that the expression of this gene is a vestigial remnant of the proposed evolutionary history of the trigger hair from an ancestral, tentacle-like secretory structure (Williams, 1976). However, we don’t see enriched expression of other genes obviously associated with volatile production. Interestingly, when we blasted the transcript against the *Arabidopsis* TAIR10 genome (https://www.arabidopsis.org/Blast/) we found a reported hit against *Arabidopsis* MSL10, due to a conserved 17 nt stretch. Thus, a shared regulatory sequence may exist between this transcript and MSL-related genes.

### Cloning of Venus flytrap cDNAs and sequencing of DmFLYC1 genomic DNA

cDNA from Venus flytrap whole-trap tissue was prepared from RNA using the Maxima First Strand cDNA Synthesis Kit (Thermo Fisher). *FLYC1* cDNA (Trinity ID # comp20014_c0_seq1) was then PCR amplified using primers CP1010 and CP1011 (**Supplemental Table 7**), which anneal in the predicted 5’ and 3’ UTRs, and was ligated into pCR-Blunt II-TOPO (Invitrogen). A clone was generated that matched the predicted sequence from the de novo transcriptome, as assessed by Sanger sequencing methods. Similarly, *EF1α* (comp10702_c0_seq1) was PCR amplified from cDNA template using primers CP1144 and CP1145, and ligated into pCR-Blunt II-TOPO. We chose this EF1α transcript as a ubiquitously expressed control gene for our RNAscope experiments due to its high expression in all RNA-seq tissue samples. We sequenced the *EF1α* cDNA insert using flanking M13F and M13R primers that anneal in the pCR-Blunt II-TOPO vector, and verified that the first ∼1.1 kb and last ∼350 bp of the cDNA matched the prediction from our *de novo* transcriptome.

Venus flytrap genomic DNA was extracted from fresh plant tissue using a CTAB-based method (Murray & Thompson, 1980). Using this as template, we PCR-amplified overlapping fragments covering the gene and sequenced these using Sanger sequencing. Overlapping and nested primer sets were: CP1010 and CP0995; CP0994 and CP1034; CP1033 and CP1035; CP1172 and CP1173; CP1176 and CP1177; and CP1174 and CP1009. The presence of two overlapping peaks on a chromatogram was used as evidence of SNPs and allelic variation. Our most 3’ primer (CP1009) anneals over the stop codon and last 31 bases of the coding sequence, and, as such, we cannot rule out the existence of additional SNPs within this stretch. To compare our sequence against that of another strain, we performed an alignment of the coding sequence against transcript Dm_00009130-RA (Palfalvi et al., 2020; https://www.biozentrum.uni-wuerzburg.de/carnivorom/resources). The three ambiguous nucleotides in Dm_00009130-RA were not included in the analysis. For electrophysiological characterization, *FLYC1, FLYC2*, and *OSCA* cDNAs were gene synthesized (human codon optimized) into pIRES2-mCherry vector from Genewiz. K558E and K579E substitutions in *FLYC1* were generated using Q5 Site-Directed Mutagenesis Kit (New England BioLabs) according to the manufacturer’s instruction and confirmed by full-length DNA Sanger sequencing.

### Identification of Drosera capensis cDNA sequences and design of qRT-PCR primers

To find *Drosera FLYC1* homologs, we blasted the first exon of Venus flytrap *FLYC1* against the scaffold assemblies of the *D. capensis* genome (Butts et al., 2016). We reasoned that because the first exon codes for the N-terminal cytoplasmic domain—which is most divergent among MSL family members—it would therefore best differentiate among different *MSL* genes and find the best *FLYC1* homolog. We found four close homologous sequences using this method (e-value of 0.0; > 60% query cover; NCBI blastn), two of which were located on scaffold LIEC01006169.1, one on LIEC01010092.1, and another on LIEC01012078.1. Position 42843-46002 of scaffold LIEC01006169.1 (reverse strand; start to stop codon) and position 23270-26408 of scaffold LIEC01012078.1 (forward strand; start to stop codon) were predicted to code for transcripts closely resembling the complete Venus flytrap *FLYC1* sequence based on inferred exon structure and conserved sequences. The intervals between the start and stop codons shared 93.7% identity, while the 3.5 kb regions immediately upstream and the 10 kb regions immediately downstream of the start and stop codons shared only 56.61% and 46.69% identity, respectively, as determined by alignment using the default settings of Clustal Omega (Sievers et al., 2011). This indicates that the two sequences are likely two different genes having arisen from an ancestral gene duplication event. Here, we call these genes *DcFLYC1*.*1* and *DcFLYC1*.*2*, respectively.

To determine the exact *DcFLYC1*.*1* and *DcFLYC1*.*2* sequences coded for in our *D. capensis* plants, the cDNAs were amplified using primers predicted to bind in the 5’ and 3’ UTRS (primers CP1240 and CP1243; and CP1224 and CP1225, respectively). Each cDNA was amplified from template derived from two different plants, and these PCR products were independently cloned into pCR-Blunt II-TOPO vector. For *DcFLYC1*.*1*, 1 of 2 and 2 of 2 clones from each of the two reactions from independent templates shared the same cDNA sequence as determined by Sanger sequencing. For *DcFLYC1*.*2*, 3 of 7 and 4 of 8 clones from each of the two reactions shared the same cDNA sequence. These sequences have been submitted to GenBank. Other clones had additional base differences not seen in any other clone. While these might represent endogenous variation in this tetraploid species, they are more likely a result of PCR amplification errors or low-fidelity transcription and reverse transcription processes.

*DcFLYC1*.*1* and *DcFLYC1*.*2* cDNAs share high sequence similarity. To resolve between the two by qRT-PCR, we designed primer pairs where one of the two primers annealed in a less-conserved stretch of residues in the 5’ or 3’ UTR (CP1242 and CP1237, and CP1224 and CP1233, respectively). The resolution of the two cDNAs using these primers was confirmed by directly sequencing the two PCR products by Sanger sequencing. *DcFLYC1*.*1* and *DcFLYC1*.*2* cDNAs were gene synthesized by Genewiz (human codon optimized) and subcloned into pIRES2-mCherry vector for electrophysiological characterization.

To design qRT-PCR primers for the *D. capensis EF1α* reference transcripts, we amplified a PCR product from cDNA template using primers CP1208 and CP1209. Sanger sequencing of this product suggested the presence of at least two highly similar *EF1α* transcripts, as evidenced by double peaks on the chromatogram. These may represent the products of different genes or alleles. To avoid biases against any one *EF1α* transcript, we refined our qRT-PCR primers to anneal over unambiguous bases (CP1218 and CP1219).

### qRT-PCR

500-1000 ng of RNA extracted from Cape sundew leaf and tentacle tissue were used to generate cDNA using the Maxima First Strand cDNA Synthesis Kit (Thermo Fisher) according to the manufacturer’s instructions. Diluted cDNA samples were then used as template for qPCR using SYBR Green/Fluorescin dye on a CFX384 Real-Time PCR Detection System (Bio-Rad). The PCR cycling condition consisted of 45 cycles of 95°C for 10 s, 60°C for 20 s, and 72°C for 30 s. Data were analyzed with Bio-Rad CFX Manager software (Version 1.6), and fold-changes in gene expression were calculated using the ΔΔCt method (Schmittgen & Livak, 2008). Comparisons were performed in biological quadruplicate.

### Bioinformatics analyses: Homology modelling, protein/nucleotide comparisons, protein topology predictions, and evolutionary history

Unless otherwise specified, all nucleotide and amino acid comparisons were made using the default settings of MUSCLE (Edgar, 2004a, 2004b). For phylogenetic tree analysis of MscS domain and OSCA proteins shown in **Figure 2A**,**D**, Maximum Likelihood trees were built from MUSCLE alignments of MscS domains (Haswell & Meyerowitz, 2006) and full-length OSCA proteins using MEGA X software (LG + G model, 4 discrete Gamma categories) and viewed using iTOL (Kumar et al., 2018; Letunic & Bork, 2019). All positions with less than 95% site coverage were eliminated (partial deletion option). The trees with the highest log likelihood and bootstrap values from 1000 replications are shown.

Protein topologies of FLYC1 and FLYC2 (**Figure. 2B**,**C**) were predicted using Protter (Omasits et al., 2014). DmOSCA topology (**Figure. 2E)** was determined by aligning the protein against *Arabidopsis* OSCA1.2 and assigning TM and pore domains at the same positions identified in the OSCA1.2 protein structure (Jojoa-Cruz et al., 2018). To model the Venus flytrap FLYC1 TM6 putative pore domain, residues A555-P593 were threaded to residues Q92-F130 of MscS in a closed conformation (PDB 2OAU) (Bass et al., 2002). C7 symmetry was imposed using Rosettascripts (Fleishman et al., 2011), and side chain and backbone conformations were minimized using the Rosetta energy function (Leaver-Fay et al., 2011) with the solvation term turned off due to exposed hydrophobics in the partial structure.

### Accession numbers

RNA-Seq data, our de novo transcriptome, ORF identification, and downstream differential gene expression analysis can be found at NCBI using the accession numbers referenced above, and under the umbrella Bioproject PRJNA530242. CDS sequences subcloned and sequenced from plant cDNA template are deposited in GenBank (*DmFLYC1, DcFLYC1*.*1*, and *DcFLYC1*.*2*).

### Cell culture and transient transfection

PIEZO1-knockout Human Embryonic Kidney 293T (HEK-P1KO) were used for all heterologous expression experiments. HEK-P1KO cells were generated using CRISPR–Cas9 nuclease genome editing technique as described previously (Lukacs et al., 2015), and were negative for mycoplasma contamination. Cells were grown in Dulbecco’s Modified Eagle Medium (DMEM) containing 4.5 mg.ml-1 glucose, 10% fetal bovine serum, 50 units.ml-1 penicillin and 50 µg.ml-1 streptomycin. Cells were plated onto 12-mm round glass poly-D-lysine coated coverslips placed in 24-well plates and transfected using lipofectamine 2000 (Invitrogen) according to the manufacturer’s instruction. All plasmids were transfected at a concentration of 700 ng.ml-1. Cells were recorded from 24 to 48 hours after transfection.

### Electrophysiology

Patch-clamp experiments in cells were performed in standard cell-attached, or excised patch (inside-out) mode using Axopatch 200B amplifier (Axon Instruments). Currents were sampled at 20 kHz and filtered at 2 kHz or 10 kHz. Leak currents before mechanical stimulations were subtracted off-line from the current traces. Voltages were not corrected for a liquid junction potential (LJP) except for ion selectivity experiments. LJP was calculated using Clampex 10.6 software. All experiments were done at room temperature.

### Solutions

For cell-attached patch clamp recordings, external solution used to zero the membrane potential consisted of (in mM) 140 KCl, 1 MgCl_2_, 10 glucose and 10 HEPES (pH 7.3 with KOH). Recording pipettes were of 1-3 MΩ resistance when filled with standard solution composed of (in mM) 130 NaCl, 5 KCl, 1 CaCl_2_, 1 MgCl_2_, 10 TEA-Cl and 10 HEPES (pH 7.3 with NaOH).

Ion selectivity experiments were performed in inside-out patch configurations. PCl/PNa was measured in extracellular solution composed of (in mM) 150 NaCl and 10 HEPES (pH 7.3 with NaOH) and intracellular solution consisted of (in mM) 30 NaCl, 10 HEPES and 225 Sucrose (pH with NaOH). Calcium-gluconate solution for Figure 4-figure supplement 2 was composed of (in mM) 50 Calcium gluconate, 0.5 CaCl_2_, 10 HEPES, 170 Sucrose (pH 7.3 with NaOH).

### Mechanical stimulation

Macroscopic stretch-activated currents were recorded in the cell-attached or excised, inside-out patch clamp configuration. Membrane patches were stimulated with 1 or 3 second negative pulses through the recording electrode using Clampex controlled pressure clamp HSPC-1 device (ALA-scientific), with inter-sweep duration of 1 minute. Pressure for half-maximal activation was calculated by fitting pressure-response curve for individual cells, which was recorded in cell-attached configuration, with Boltzman equation and averaging P_50_ values across all cells. Stretch-activated single-channel currents were recorded in the cell-attached configuration. Since single-channel amplitude is independent of the pressure intensity, the most optimal pressure stimulation was used to elicit responses that allowed single-channel amplitude measurements. These stimulation values were largely dependent on the number of channels in a given patch of the recording cell. Single-channel amplitude at a given potential was measured from trace histograms of 5 to 10 repeated recordings. Histograms were fitted with Gaussian equations using Clampfit 10.6 software. Single-channel slope conductance for each individual cell was calculated from linear regression curve fit to single-channel I-V plots.

### Permeability ratio measurements

Reversal potential for each cell in the mentioned solution was determined by interpolation of the respective current-voltage data. Permeability ratios were calculated by using the following Goldman-Hodgkin-Katz (GHK) equations:

PCl/PNa ratios:

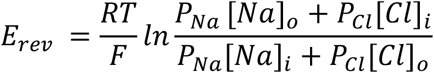

### In situ hybridization and imaging

Whole Venus flytrap and Cape sundew leaves were cut from the plant, fresh frozen in liquid N2, and 15 µm sections collected. RNA in situ hybridization (RNAscope) was performed on sections according to manufacturer’s instructions (ACDBio: #323100) using probes against *DmFLYC1* (ACDBio; Ref: 546471, lot: 18177B), *DmFLYC1*-*sense* (ACDBio; Ref: 566181-C2, lot: 18361A), *DmFLYC2* (ACDBio; Ref: 546481, lot: 18177C), *DmOSCA* (ACDBio; Ref: 571691, lot: 19032A), *DcFLYC1*.*1/DcFLYC1*.*2* (ACDBio; Ref: 572451, lot: 19037B), and *EF1α* (ACDBio; Ref: 559911-C2, lot: 18311B). *DmFLYC1-sense* (**Figure 3B**) and *DmFLYC2* (**Figure 3-figure supplement 2)** were tested on the same section. However, *DmFLYC2* probe was independently tested in two additional experiments and no signal was observed. Slides were mounted with Vectashield + DAPI (Ref: H1200, lot: ZE0815). Stained sections were imaged with a Nikon C2 laser scanning confocal microscope and z-stacks were acquired through the entire section with a 60x objective. Displayed images are max projections of the entire z-stack. Images were processed using ImageJ (Fiji image processing package).

