## Supplemental Figures for "Stretch-activated ion channels identified in the touch-sensitive structures of carnivorous Droseraceae plants"

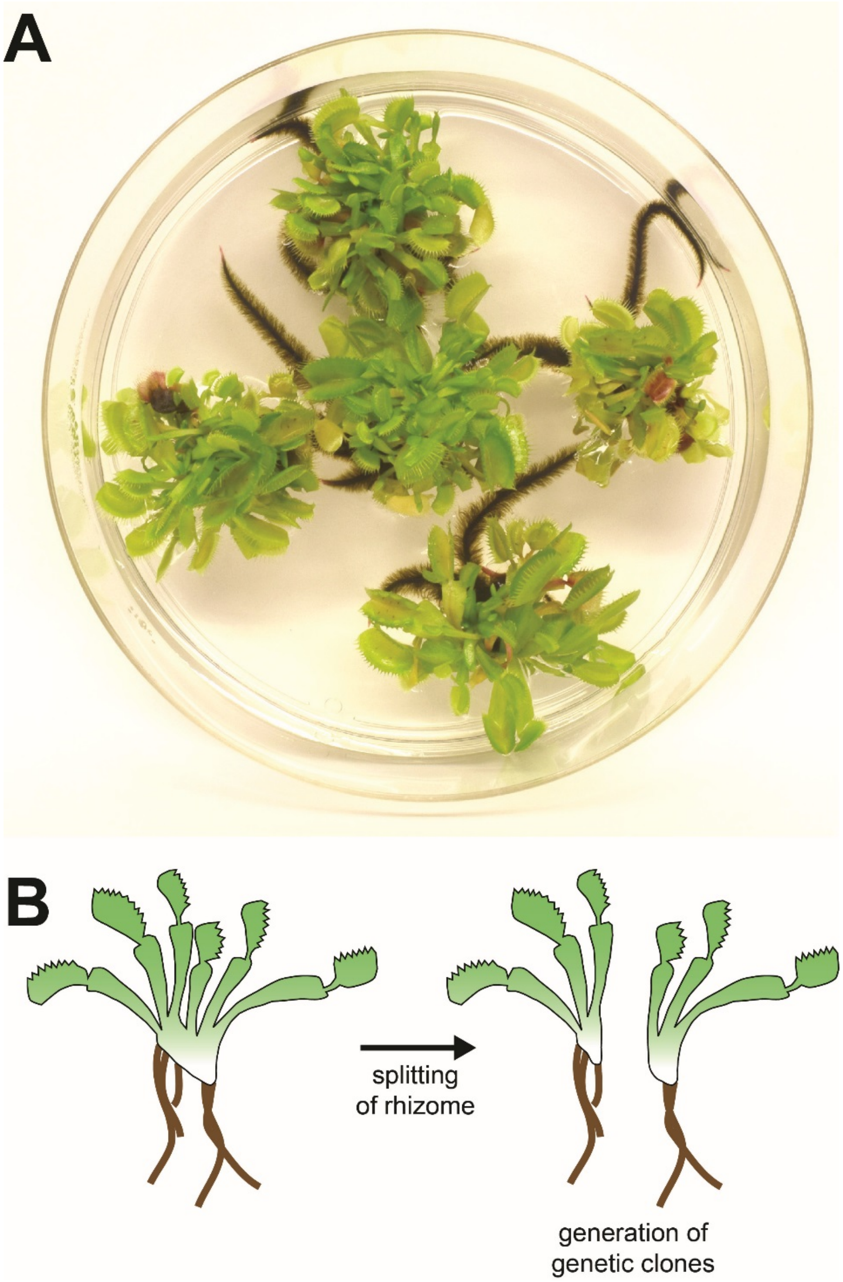


**Figure 1- figure supplement 1. Venus flytrap clonal propagation system.** (**A**) Example of clonal Venus flytraps growing in tissue culture (10 cm plate) using methods adapted from Jang et al., 2003. (**B**) Diagram depicting the method of propagation. Rosettes were separated by splitting the rhizome. Plants were then transferred to fresh sterile growth medium for further propagation in culture or transferred to soil and 'hardened' for at least 2-3 months prior to experiments.

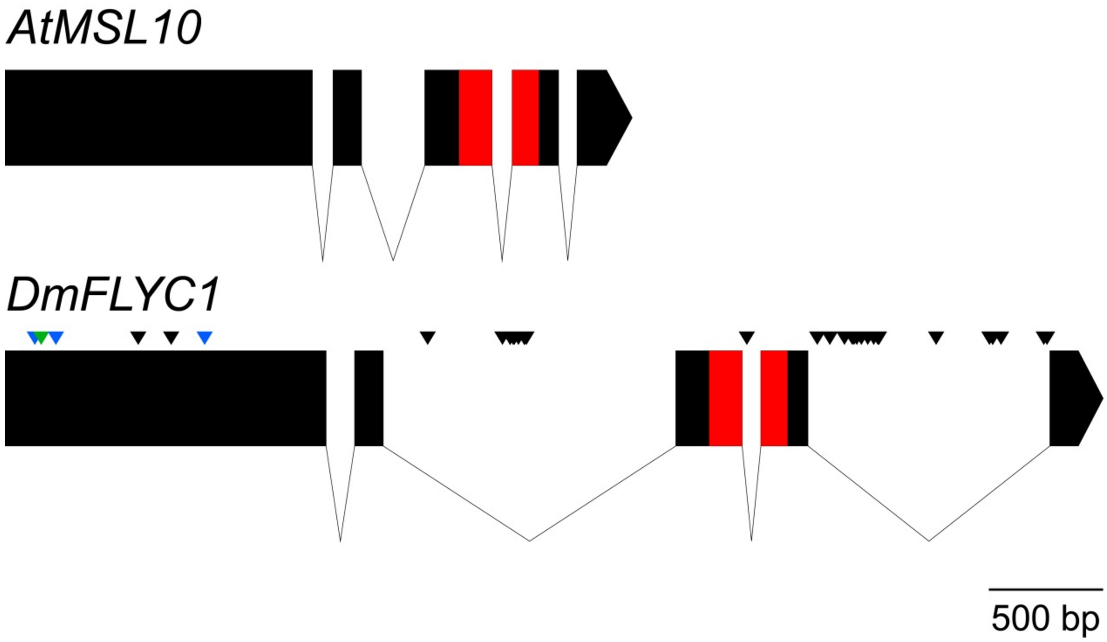


**Figure 1- figure supplement 2. Gene structure of Venus flytrap *FLYC1*.** A comparison of *Arabidopsis MSL10* and Venus flytrap *FLYC1* genes. Boxes represent exons; lines, introns. The sequence between the start and stop codons only is shown. SNPs (allelic differences) in *FLYC1* are indicated for our strain by black arrowheads. 32 SNPs in total were detected, of which only 2 were found in the coding region and were both silent. When the coding sequence (exons only) was compared to that of another strain (Palfalvi et al, 2020), an additional four SNPs were identified, three of which were silent and one which caused the amino acid change A51G relative to our strain (indicated by blue and green arrowheads, respectively).

**Figure 1- Supplemental Table 1. Size estimates of two *Arabidopsis* (Col-0) samples compared to our Venus flytrap strain (CP01).**

| Species | DNA Content (pg/2C) | St. Dev. |
|---|---|---|
| ***A. thaliana* (Col-0) Sample #1** | 0.38 | 0.010 |
| ***A. thaliana* (Col-0) Sample #2** | 0.40 | 0.007 |
| **D. muscipula (CP01)** | 7.86 | 0.359 |

**Figure 1- Supplemental Table 6. Summary of sequencing reads used to build the *de novo* transcriptome (NCBI Transcriptome Shotgun Assembly Sequence Database accession # GHJF00000000).**

| Sample | Paired replicates | Tissue | # paired reads | SRA accession # |
|---|---|---|---|---|
| unfed1 | no | 2 traps | 9,079,584 | SRR8834216 |
| unfed2 | no | 2 traps | 9,424,911 | SRR8834215 |
| fed1 | no | 2 traps | 9,597,638 | SRR8834218 |
| fed2 | no | 2 traps | 9,887,029 | SRR8834217 |
| trap1 | yes (A) | ~20 traps[a] | 4,410,249 | SRR8834220 |
| trigger_hair1 | yes (A) | 250 hairs | 3,702,412 | SRR8834221 |
| trap2 | yes (B) | ~20 traps[a] | 10,974,314 | SRR8834219 |
| trigger_hair2 | yes (B) | 750 hairs | 9,711,759 | SRR8834214 |
| trap3 | yes (C) | ~20 traps[a] | 11,418,472 | SRR8834222 |
| trigger_hair3 | yes (C) | 750 hairs | 9,975,164 | SRR8834213 |

[a]Trigger hairs removed.

```
FLYC1        -------MGSYLHEPPGDEP-------------------------SMRIEQPKTADRAPEQ   29
FLYC2        MEGVRNPLRNSFNKAHEAEPQRKKNLEQEERLILLQHRNDPNSQSFSSEDPNSLLLQVKV   60

FLYC1        VAIHICEPSKVVTE----------------------------------------SFPFSET   50
FLYC2        EVAGSCDPAKTAVPTKPPVSPGGGGNLIWRDSSYDFRNDVVKGCSRDTDDDSGEFDFQKH  120

FLYC1        AEPEA-KSKNCPCPEIARIGPCPNKPPKIP--INRGLSRISTNKSRPKSRFGEPSWPVES  107
FLYC2        RVAEEEDEGEERDPESQTLSPVSESPHEYGKITPRGAAKVSFKESELVHR-----RPSDG  175

FLYC1        SLDLTSQSPVSPYREEAFS----VENCGT-------------------AGSRRGSFARGT  144
FLYC2        GVF-AADGVSASVRDEEVVMCTSNASCQRKSTSTRVKTKSRLLDPPDDGDTRSGRILRSG  234

                                                                   TM1
FLYC1        TSRAASSSRKDETKEGPDEKEVYQRVTAQLSARNQKRMTVKLMIELSVFLCLLGCLVCSL  204
FLYC2        LMPRSEDHEDEDPFSGEDIPEEYKKM-----------KFSFLSAVELVSLLLIIAGLVCSV  284

                                         TM2
FLYC1        TVDGFKRYTVIGLDIWKWFLLLLVIFSGMLITHWIVHVAVFFVEWKFLMRKNVLYFTHGL  264
FLYC2        VIPVVRRVTVWDMQLWKWEVMVLVLICGGLVSGWLIRFVVFFIERNFLLRKRVLYFVYGL  344

                        TM3                            TM4
FLYC1        KTSVEVFIWITVVLATWVMLIKPDVNQPHQTRKILEFVTWTIVTVLIGAFLWLVKTTLLK  324
FLYC2        RRAVQRCLWLGWVLIAWRLILDKKVEKETNSRS-LLYVTKILVCLVVGTLIWLLKTLLVK  403

FLYC1        ILASSFHLNRFFDRIQESVFHHSVLQTLAGRPVVELAQGISRTES---------------  369
FLYC2        VLAMSFHVSTFFDRIQEALFDQYVIETLSGPPTIEIQHVKEDEDQVMLEVQKLQSAGLSI  463

FLYC1        -------------------------QDGAGQV-------------SFMEHTKTQNKKVVDV  392
FLYC2        PAELKATCLPNVNVNGKPVGSDPGPTPGVGKSPRSGVIGKSPRFSRAMPEKEEGAGGITI  523

FLYC1        GKLHQMKQEKVPAWTMQLLVDVVSNSGLSTMSGMLDEDMVEGGVELD-DDEITNEEQAIA  451
FLYC2        DHLHRLNQKNISAWNMKRLMNIVRYGVLSTL----DEQILESGIEDEPSLHIKNENQAKA  579

FLYC1        TAVRIFDNIVQDKVDQSYIDRVDLHRFLIWEEVDHLFP-LFEVNEKGQISLKAFAKWVVK  510
FLYC2        AAKRLFKNVARPGSKCIYLE--DLMRFMREDEAARTMRAIEGSAESKGISKIALKNWVVN  637

                                                      TM5
FLYC1        VYNDQAALKHALNDNKTAVKQLNKLVTAILIVMMIVIWLIVTGIATTKLIVLLSSQLVVA  570
FLYC2        VFRERRALALSLNDTKTAVNKLHQLLNFIVGFTIAIIWLLILGVPMTHFFVFITSQLLLL  697

                TM6
FLYC1        AFIFGNTCKTIFEAIIFVFVMHPFDVGDRCVIDGNKMLVEEMNILTTVFLKWDKEKVYYP  630
FLYC2        TFMFGNTFKTTFEAIIFLFVMHPFDVGDRCEVEGVQMIVEEMNILTTVFLRYDNLKITYP  757

FLYC1        NSILCTKAIGNFFRSPDQGDVLEFSVDFTTPVLKIGDLKDRIKMYLEQNLNFWHPQHNMV  690
FLYC2        NSVLATKPINNYYRSPEMGDSVDFCVHISTPVEKIVVMKERITRYMESRRDHWRPSPKVV  817

FLYC1        VKEIENVNKIKMALFVNHTINFQDFAEKNRRRSELVLELKKIFEELDIKYNLLPQEISIR  750
FLYC2        MREVEDMNRLKFSVWMCHTMNHQDMGERWARRELLVVEMVKIFKELDVQYRMLPHDVNVR  877

FLYC1        NM------------------   752
FLYC2        TMPSLVCDRLPSNWITCTGK   897
```

**Figure 2- figure supplement 1. Sequence alignment of DmFLYC1 and DmFLYC2 proteins.** Conserved amino acids are shown in red. Similar and highly similar amino acids are in blue and orange, respectively. For FLYC1, also shown are the transmembrane (TM) domains shadowed in gray as predicted by Protter (Omasits et al., 2014), and the residues forming the putative pore domain modeled in Figure 4- figure supplement 3 are boxed.

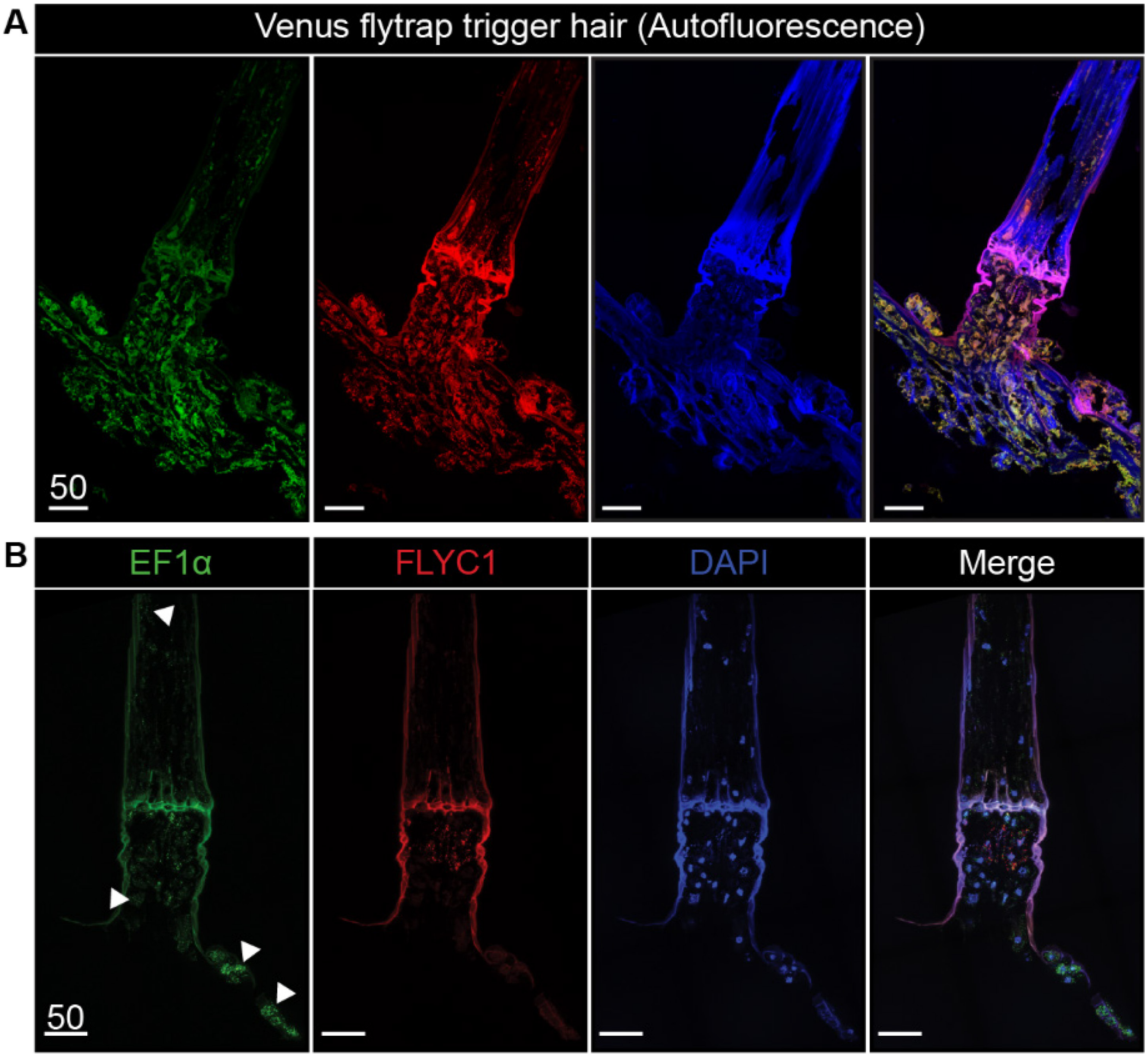


**Figure 3- figure supplement 1. Autofluorescence and *EF1α* transcript expression in Venus flytrap.** (**A**) Unprocessed Venus flytrap trigger hair in different channels to depict autofluorescence from the plant cuticle. Images were taken with the same settings as tissue subjected to *in situ* hybridization. (**B**) A longitudinal section of a trigger hair and trap with smFISH against the housekeeping gene *EF1α* (green) and *FLYC1* (red) as well as DAPI (blue). White arrowheads indicate examples of *EF1α* expression in various cells throughout the tissue. Scale bar, µm.

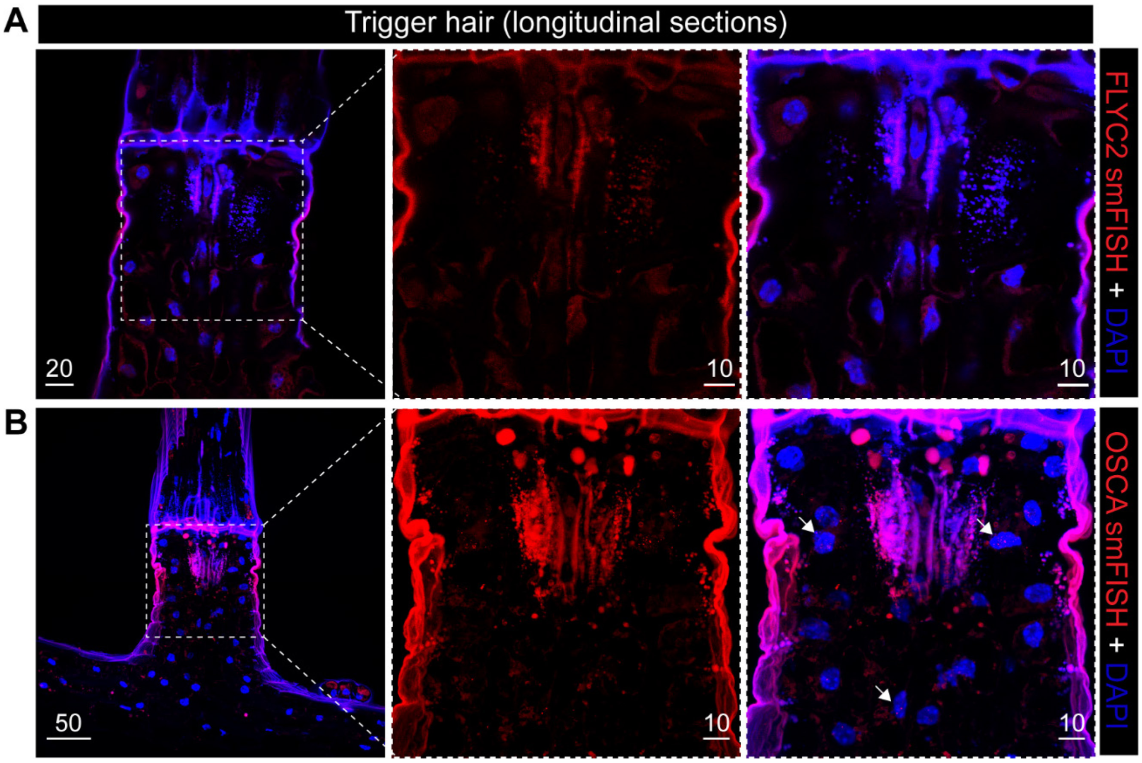


**Figure 3- figure supplement 2. FLYC2 and DmOSCA transcript expression in trigger hair.** Max projection through longitudinal sections after fluorescent *in situ* hybridization of (**A**) *DmFLYC2* transcript and (**B**) *DmOSCA* transcript at low (left) and high resolution (center and right). *DmFLYC2* and *OSCA* transcript is depicted in red channel and DAPI in blue. Note no transcript was observed for *DmFLYC2*, whereas *OSCA* was observed mostly in the sensory cells. Scale bars, µm.

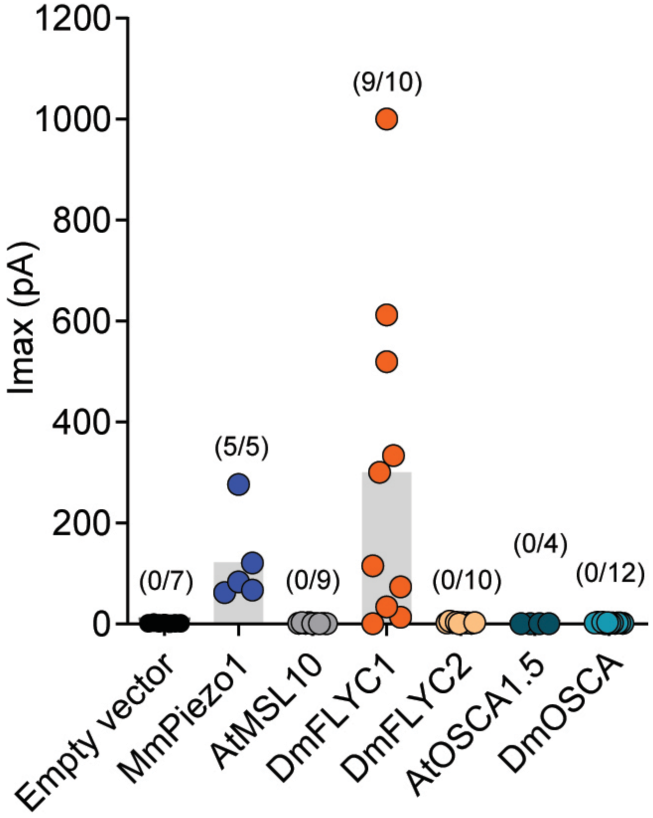


**Figure 4- figure supplement 1. DmFLYC2 and DmOSCA functionality.** Macroscopic stretch-activated currents recorded from HEKP1-KO cells transfected with Mock (N=7), MmPiezo1 (N=5), AtMSL10 (N=9), DmFLYC1 (N=10), DmFLCY2 (N=10), AtOSCA1.5 (N=4), and DmOSCA (N=12) plasmids. Empty vector, MmPiezo1, and DmFLYC1 data is the same as Figure 4A.

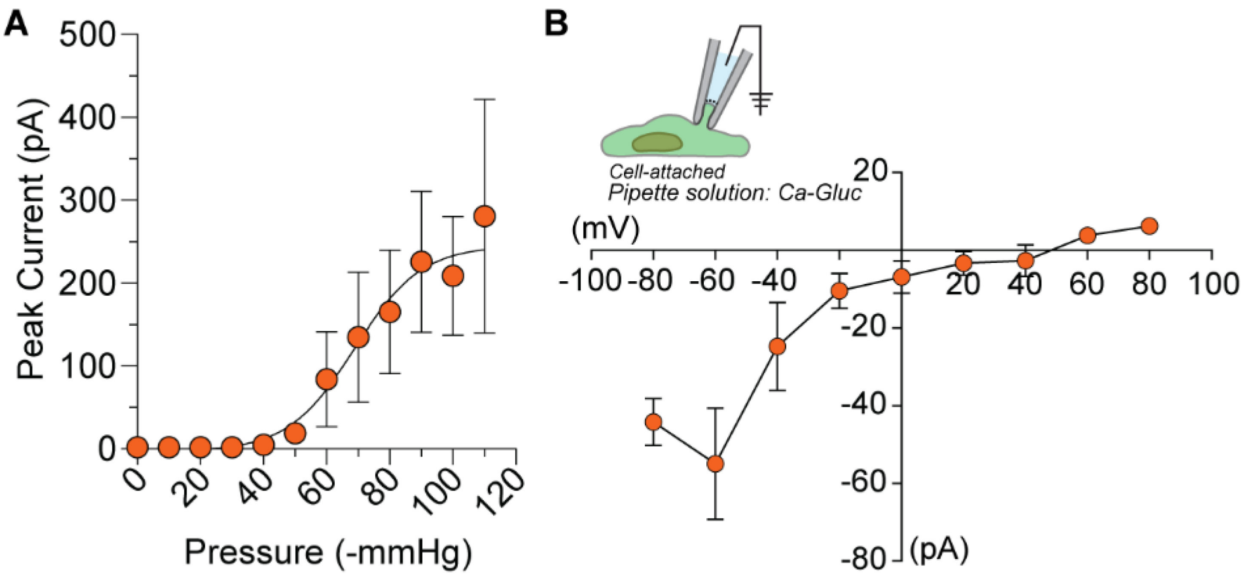


**Figure 4- figure supplement 2**. **Half maximal pressure response and chloride permeability in DmFLYC1.** (**A**) Average pressure-response curve across different cells (N=9). Same data as Figure 4B but the peak current is not normalized to maximal response. (**B**) Average I-V (N=5) of DmFLYC1 stretch-activated currents recorded in cell attached patch clamp configuration. The recording pipette is composed of (in mM) 50 Calcium gluconate, 0.5 $CaCl_2$, 10 HEPES, 170 Sucrose (pH 7.3).

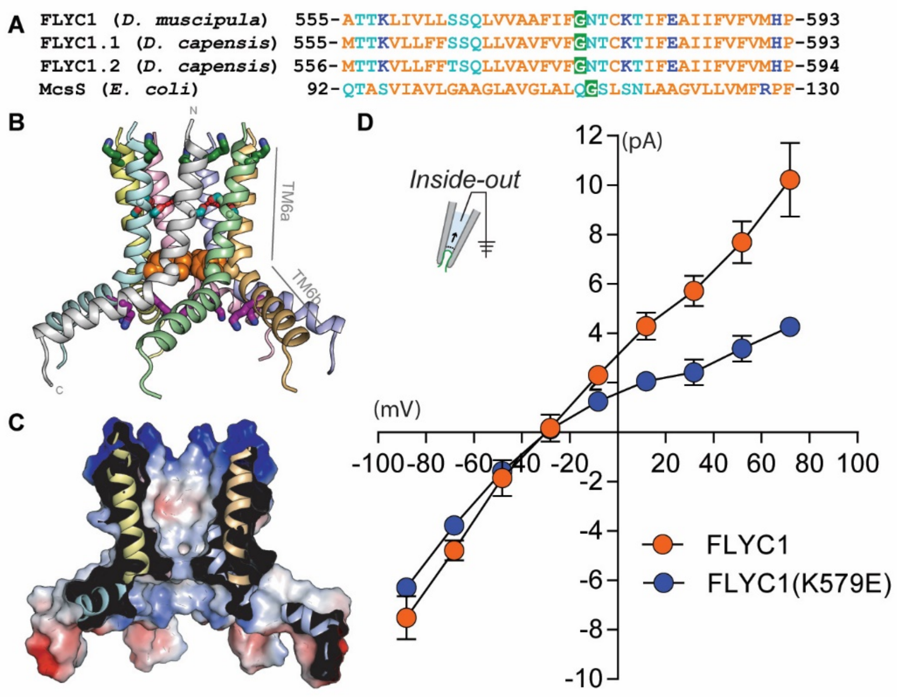


**Figure 4- figure supplement 3. The sequence of a putative pore-forming helix in FLYC1 is compatible with MscS-like channel structure.** (**A**) Sequence alignment of the putative pore helix of Venus flytrap FLYC1 and *Drosera* DcFLYC1.1 and DcFLYC1.2 proteins with MSL10 and MscS. Nonpolar, polar, and ionizable residues are orange, light blue, and dark blue, respectively. A glycine predicted to localize at a central bend in the helix is shaded green. (**B**) Modeled heptameric organization. The sequence of Venus flytrap FLYC1 was threaded on the inner helix of heptameric MscS in a closed conformation (PDB 2OAU), and minimized using the Rosetta energy function while imposing $C_7$ symmetry. Subunits are shown in different colors. Basic residues K558 and K579 are green and purple sticks, respectively. Predicted intersubunit hydrogen bonds between serines S564 and S565 (blue sticks) of pore segments are shown with red dashes, while a ring of phenylalanines (F572; orange spheres) constrict the pore. Helices TM6a, forming the central pore, and amphipathic helix TM6b are indicated for one pore segment. (**C**) Cross section through the protein surface colored by electrostatic potential, showing an uncharged (white) pore, with positive charge (blue) above and below the pore. (**D**) Average I-V of stretch-activated single channel currents from FLYC1 (N=7) and FLYC1(K579E) (N=4) in asymmetrical NaCl solution. FLYC1 data is the same as Fig 4D. Extracellular solution (pipette solution) composition was (in mM) 150 NaCl and 10 HEPES (pH 7.3) and intracellular solution (bath solution) composition was 30 NaCl, 10 HEPES and 225 Sucrose (pH 7.3).

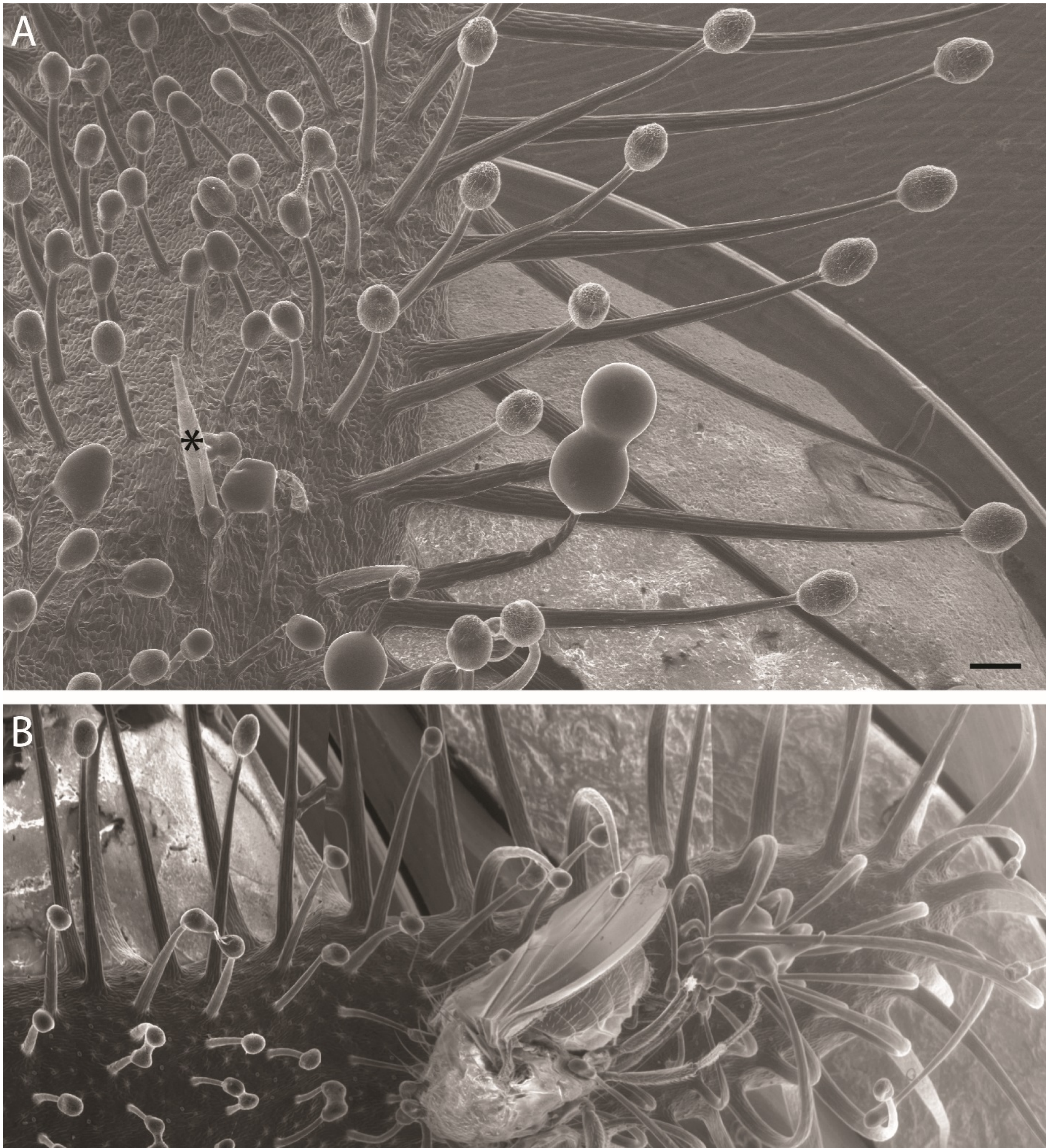


**Figure 5- figure supplement 1. Variation in Cape sundew tentacles.** (**A**) Scanning electron microscopy (SEM) image showing mucilage-producing tentacles of different lengths on the Cape sundew leaf blade. An asterisk (*) marks the staging needle. Tentacles increase in length towards the leaf edge. (**B**) A composite image of three SEM micrographs showing tentacle bending around a *D. melanogaster* fly. Scale bars, 200 µm.

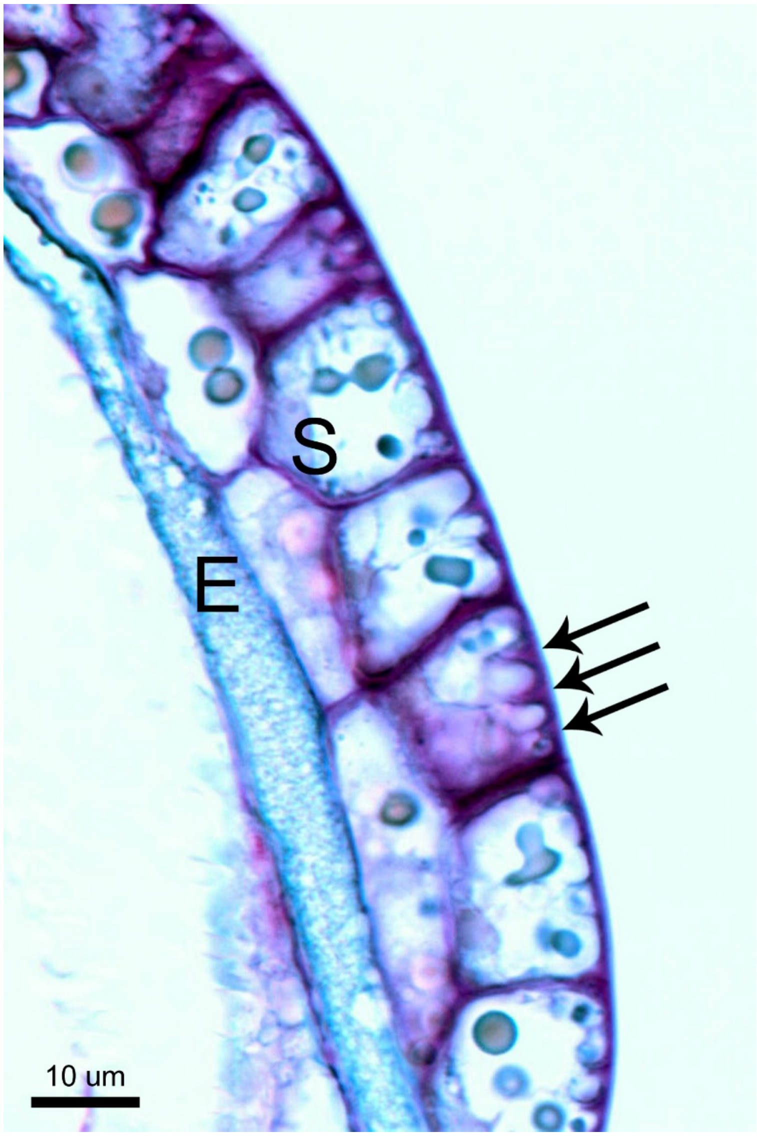


**Figure 5- figure supplement 2. Unique cell morphology of Cape sundew sensory/excretory cells.** Toluidine blue-stained longitudinal section through the head of a Cape sundew tentacle. E, endodermis-like cells; S, outer two layers of secretory cells. Arrows mark the unique cell wall buttresses of a single outer secretory cell, around which membrane crenellations occur. See also Lloyd, 1942.

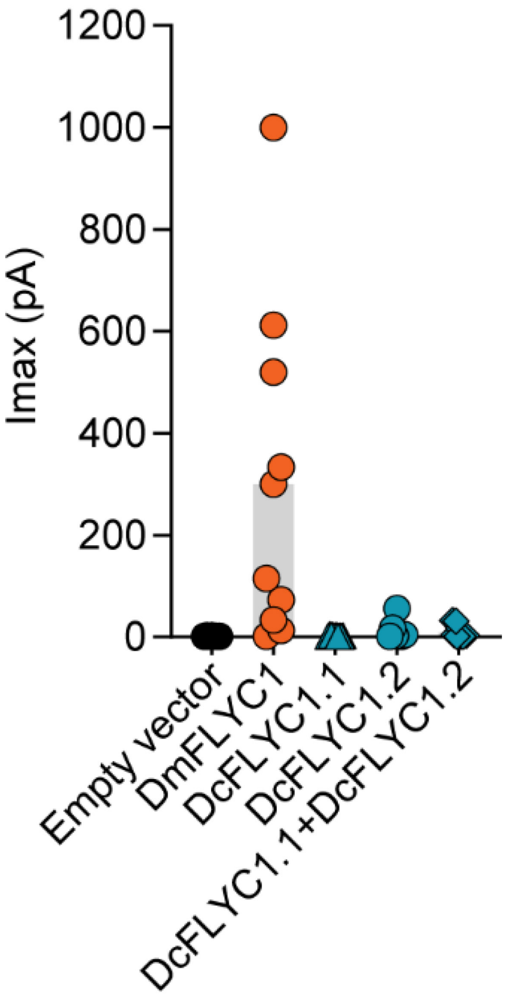


**Figure 5-figure supplement 3. Drosera FLYC functionality.** Macroscopic stretch-activated currents recorded from HEKP1-KO cells transfected with Mock (N=7), DmFLYC1 (N=10), DcFLCY1.1 (N=7), DcFLYC1.2 (N=10), and DcFLYC1.1/1.2 (N=11) plasmids. Empty vector, and DmFLYC1 data is the same as Figure 4A.

**Methods. Primer sequences.**

| Primer | Sequence (5’ → 3’) |
|---|---|
| CP0994 | CCAGTGTCACCTTATAGGGAAGAAGCG |
| CP0995 | CCCTCGACGTAGTTCCCCTAGC |
| CP1009 | cacatcgatTCACATATTGCGGATACTAATTTCTTGGGGC |
| CP1010 | acacccgggGCTAGCTTTCATCCACCGAATAAACACC |
| CP1011 | cacatcgatACATCATTGACCAGAAGCAAGGCACTC |
| CP1033 | TGGCATCTTCATTCCATTTGAATAGGTTCTTTG |
| CP1034 | GCATACCCGACATGGTCGACAAGC |
| CP1035 | TGACAAGTTTGTTTAACTGCTTCACTGCTG |
| CP1144 | AGGTCTTTAGATTAACTCTTCAACATGGGTAAGG |
| CP1145 | ACAAGACTTCATTTTGCACCCTTCTTTATCG |
| CP1172 | GATTGAGCAAACAAAAGGCGCATGAAG |
| CP1173 | AAATTTTGACCTACGTTGACCGTCAGC |
| CP1174 | TGATCAGGCTGCGCTTAAACATGC |
| CP1176 | ACATTCTACTTTGTTTGCAATTGTTTTCCCACTC |
| CP1177 | AAGAAACATTAAGCTGCACCTGCTCC |
| CP1208 | GAGAGGTCCACCAACCTTGACTGG |
| CP1209 | AGCAACGGTCTGACGCATGTCC |
| CP1218 | GTGCCAGTGGGAAGAGTTGAGAC |
| CP1219 | CAGAGAAAGTCTCGACAACCATGGG |
| CP1224 | CAAATCATTGAAAACGTGAAGGGAAGCACTG |
| CP1225 | CCTATGAGATACTTAGCCTGTTAGCCATGC |
| CP1233 | GCCCTTCTATAGTAGTCTCACCTCTTCG |
| CP1237 | GCCATGCGCAGCATATGTACTAGC |
| CP1240 | GCTATTTCTTATTCTCCTGAGCACAACATACTG |
| CP1242 | TGATCGCTGTGTCGTAGATGGAACAATG |
| CP1243 | CTAATGGATTGCAAACTAGGAGATGCTTAGC |

**Supplementary Video 1. Touch response of the Venus flytrap.**
Real time recording of trap closure in response to insect (house fly) touch.

**Supplementary Videos 2 and 3. Touch response of the Cape sundew plant.**
Time lapse recordings of tentacle bending in response to insect (*Drosophila melanogaster*) touch. Insects were placed directly on the leaf tentacles and recordings started within 10 seconds thereafter. 1 second represents 5 min.

**Protein sequences:**

**DmFLYC1**
MGSYLHEPPGDEPSMRIEQPKTADRAPEQVAIHICEPSKVVTESFPFSETAEPEAKSKNCPCPE
IARIGPCPNKPPKIPINRGLSRISTNKSRPKSRFGEPSWPVESSLDLTSQSPVSPYREEAFSVE
NCGTAGSRRGSFARGTTSRAASSSRKDETKEGPDEKEVYQRVTAQLSARNQKRMTVKLMIELSV
FLCLLGCLVCSLTVDGFKRYTVIGLDIWKWFLLLLVIFSGMLITHWIVHVAVFFVEWKFLMRKN
VLYFTHGLKTSVEVFIWITVVLATWVMLIKPDVNQPHQTRKILEFVTWTIVTVLIGAFLWLVKT
TLLKILASSFHLNRFFDRIQESVFHHSVLQTLAGRPVVELAQGISRTESQDGAGQVSFMEHTKT
QNKKVVDVGKLHQMKQEKVPAWTMQLLVDVVSNSGLSTMSGMLDEDMVEGGVELDDDEITNEEQ
AIATAVRIFDNIVQDKVDQSYIDRVDLHRFLIWEEVDHLFPLFEVNEKGQISLKAFAKWVVKVY
NDQAALKHALNDNKTAVKQLNKLVTAILIVMMIVIWLIVTGIATTKLIVLLSSQLVVAAFIFGN
TCKTIFEAIIFVFVMHPFDVGDRCVIDGNKMLVEEMNILTTVFLKWDKEKVYYPNSILCTKAIG
NFFRSPDQGDVLEFSVDFTTPVLKIGDLKDRIKMYLEQNLNFWHPQHNMVVKEIENVNKIKMAL
FVNHTINFQDFAEKNRRRSELVLELKKIFEELDIKYNLLPQEISIRNM*

**DmFLYC2**
MEGVRNPLRNSFNKAHEAEPQRKKNLEQEERLILLQHRNDPNSQSFSSEDPNSLLLQVKVEVAG
SCDPAKTAVPTKPPVSPGGGGNLIWRDSSYDFRNDVVKGCSRDTDDDSGEFDFQKHRVAEEEDE
GEERDPESQTLSPVSESPHEYGKITPRGAAKVSFKESELVHRRPSDGGVFAADGVSASVRDEEV
VMCTSNASCQRKSTSTRVKTKSRLLDPPDDGDTRSGRILRSGLMPRSEDHEDEDPFSGEDIPEE
YKKMKFSFLSAVELVSLLLIIAGLVCSVVIPVVRRVTVWDMQLWKWEVMVLVLICGGLVSGWLI
RFVVFFIERNFLLRKRVLYFVYGLRRAVQRCLWLGWVLIAWRLILDKKVEKETNSRSLLYVTKI
LVCLVVGTLIWLLKTLLVKVLAMSFHVSTFFDRIQEALFDQYVIETLSGPPTIEIQHVKEDEDQ
VMLEVQKLQSAGLSIPAELKATCLPNVNVNGKPVGSDPGPTPGVGKSPRSGVIGKSPRFSRAMP
EKEEGAGGITIDHLHRLNQKNISAWNMKRLMNIVRYGVLSTLDEQILESGIEDEPSLHIKNENQ
AKAAAKRLFKNVARPGSKCIYLEDLMRFMREDEAARTMRAIEGSAESKGISKIALKNWVVNVFR
ERRALALSLNDTKTAVNKLHQLLNFIVGFTIAIIWLLILGVPMTHFFVFITSQLLLLTFMFGNT
FKTTFEAIIFLFVMHPFDVGDRCEVEGVQMIVEEMNILTTVFLRYDNLKITYPNSVLATKPINN
YYRSPEMGDSVDFCVHISTPVEKIVVMKERITRYMESRRDHWRPSPKVVMREVEDMNRLKFSVW
MCHTMNHQDMGERWARRELLVVEMVKIFKELDVQYRMLPHDVNVRTMPSLVCDRLPSNWITCTG
K*

**DmOSCA**
MESNPEYIASLGDIVVAAVINIFFAFVFFIAFAIFRIQPVNDRVYYTKWYLRGLRSSSTNPDAF
VRKCVNLSFGSYLKFLNWMPAALQMPETELIQHAGLDSAVYLRIYLVGLKIFIPITILALSIVI
PVNWTDGGLEKSKLIAFNNLDKLSISNIRPGSEKFWTHIGMAYTVTFWACYILKKEYESIESMR
LQFLASSGRKPEQFTVLVRNVPLDSDESTSELVEHFFKVNHPDDYLTRQVIYDANVLTDLVRER
KKKQMWLNFYQLKYTRSQSRKPFCKTGFLGLWGTKVDAIDYYTMEVERLSKEISSKREMIANDT
KAVMLAAFVSFKTRRGAAICAHTQQARNPTLWLTQWAPEPRDIYWRNLAIPYASLSIRKLIVSV
TFFFLATFFMIPIAFVQSLANIEGIEKALPFLRPVIEARFVKSIIQGFLPGIVLKIFLTFLPSI
LMMMCKSEGIISLSALERRAAARYYVFLLINVFLGSIVTGTAFEQLNNILHETANAIPETIGAA
IPMKVTFFITYTMVDGWAGMAAEILRLKPLICYHLKVCFLVNTEKDKEEAMNPQSFGFNTREPQ
IQLYFLVALVYAVAAPILLPFIVLLFSLGYIVYRHQIINVYNQEYESGAAFWPDVHKRIVVALV
VSQLLLLGLLSTKKASHSTPLLVALPVLTISFHYLCKGRFLPAFVTHPLQEATLKDSMDLAREP
GLHFKRYLQNAYTHPLLKVGDNAETDEAFQEVEQGCQLVQTKRQLWRTFS*
**DcFLYC1.1**

MASNTNISQQGGEINFEKQMAHRRRHEQLAIQIPVKTASQTFRFNEEVDTRSKFSPAPDITMFY
PQPSPNKPPRVPNRTLTRRSTTLKTKPKSRFGEPSLPIDPAALWELAPNSPTPSFREATPSSNN
HRFSVGRGSSFAKGVTPRVAASSQRGETTIEGPDEKEVYERVTAQLSARDKKRMTVKLLIELAI
FLFVSGCLISSLTIHGLKVRKIYGLPIWRLFLFLLVILSGMLVTHWMIHVVVFLIEWKFLLKKN
VVYFTHGLKTSVEVFIWITLILATWGLLIEPDVRHTNRIRNALDFITWTLLSLLLGSFLWLIKT
IMIKTLAASFHLNRFFDRIQESIFHHYVLQTLSGRPVVELASGVLTRTETHNGMVSFTEHTKTH
KEKKMVDMGKLHQMKQEKVPDWTMQLLVDVVSNSGLSTMSGILDEDMAEGGVELDDDEITSEEQ
AIATAVRIFYNIVKDKDDQSYIDRKDLHRFLICEEVDLVFPLFEVKDKDQINLKAFSKWVVKLF
KERQALKHALNDNKTAVKQLDKLVTSILIVVIIAVWLLLTEIMTTKVLLFFSSQLLVAVFVFGN
TCKTIFEAIIFVFVMHPFDVGDRCVVDGTMMLVEEMNILTTVFLKWDKEKVYYPNAVLSTKAIG
NYYRSPDQVDSLEFSIDFRTPLSKIGEIKERIKKYLHQNPHLWHPNHNFVVKEIENVNKIKMQL
IFNHTINFQEFPERMKRRSELVLELKKIFEELDIKYNLLPQEVILNKVSP*

**DcFLYC1.2**

MASNTNISQQGGEINFEKQMAHRRRHEQLAIQIPVKTASQTFPFNEEVDTTRSKFSPAPDITMF
YPQPSPNKPPRVPNRNLSRRSTTLKTKPKSRFGEPSLPIDPAALWELAPNSPAPSFREATPSSN
NHRASVGRGSSFVKGVTPRVAASSRRGETTIEGPDEREVYERVTAQLSARDKKRMTVKLLIELA
VFLFVSGCLISSLTIHGLKVRIICGLPIWRLFLFLLVILSGMLVTHWMLHVVVFLIEWKFLLKK
NVVYFTHGLKTSVEVFIWITLILATWALLIEPDVRHTNRIRNALDFITWTLLSLLLCSFLWLIK
TIMIKTLAASFHLNRFFDRIQESIFHHYVLQTLSGRPVVELASGVLTRTETHNGMVSFTEHTKT
HTEKKMVDMGKLHQMKQEKVPDWTMQLLVDVVSNSGLSTMSGILDEDMAEGGVELDDDEITSEE
QAIATAVRIFYNIVKDKDDQTYIDRKDLHRFLICEEVDLVFPLFEVKDKDQISLKAFSKWVVKL
FKERQALKHALNDNKTAVKQLDKLVTSILIVVIIAVWLLLTELMTTKVLLFFTSQLLVAVFVFG
NTCKTIFEAIIFVFVMHPFDVGDRCVIDGTTMLVEEMNILTTVFLKWDKEKVYYPNAVLSTKAI
GNYYRSPDQVDSLEFSIDFRTPLSKIGEIKERIKKYLHQNPHLWHPNHNLVVKEIENVNKIKTQ
LIFNHTMNFQEFPERMKRRTELVLELKKIFEELDIKYNLLPQEVILNNVGP*
